# Combination and competition between path integration and landmark navigation in the estimation of heading direction

**DOI:** 10.1101/2021.06.30.450548

**Authors:** Sevan K. Harootonian, Arne D. Ekstrom, Robert C. Wilson

**Affiliations:** Department of Psychology, Princeton University, Princeton, NJ, USA; Department of Psychology, University of Arizona, Tucson, AZ, USA; McKnight Brain Institute, University of Arizona, Tucson, AZ, USA; Cognitive Science Program, University of Arizona, Tucson, AZ, USA

## Abstract

Successful navigation requires the ability to compute one’s location and heading from incoming multisensory information. Previous work has shown that this multisensory input comes in two forms: body-based idiothetic cues, from one’s own rotations and translations, and visual allothetic cues, from the environment (usually visual landmarks). However, exactly how these two streams of information are integrated is unclear, with some models suggesting the body-based idiothetic and visual allothetic cues are combined, while others suggest they compete. In this paper we investigated the integration of body-based idiothetic and visual allothetic cues in the computation of heading using virtual reality. In our experiment, participants performed a series of body turns of up to 360 degrees in the dark with only a brief flash (300ms) of visual feedback *en route*. Because the environment was virtual, we had full control over the visual feedback and were able to vary the offset between this feedback and the true heading angle. By measuring the effect of the feedback offset on the angle participants turned, we were able to determine the extent to which they incorporated visual feedback as a function of the offset error. By further modeling this behavior we were able to quantify the computations people used. While there were considerable individual differences in *performance* on our task, with some participants mostly ignoring the visual feedback and others relying on it almost entirely, our modeling results suggest that almost all participants used the same *strategy* in which idiothetic and allothetic cues are combined when the mismatch between them is small, but compete when the mismatch is large. These findings suggest that participants update their estimate of heading using a hybrid strategy that mixes the combination and competition of cues.

**Author summary:** Successful navigation requires us to combine visual information about our environment with body-based cues about our own rotations and translations. In this work we investigated how these disparate sources of information work together to compute an estimate of heading. Using a novel virtual reality task we measured how humans integrate visual and body-based cues when there is mismatch between them — that is, when the estimate of heading from visual information is different from body-based cues. By building computational models of different strategies, we reveal that humans use a hybrid strategy for integrating visual and body-based cues — combining them when the mismatch between them is small and picking one or the other when the mismatch is large.

## Introduction

The ability to navigate — to food, to water, to breeding grounds, or even to work — is essential for survival in many species. To navigate effectively we need to continuously update our estimates of location and heading in the environment from incoming multisensory information [1–3]. This multisensory input comes in two forms: body-based idiothetic cues, from one’s own rotations and translations (in humans generated from the vestibular, proprioceptive, and motor efferent copy systems), and visual allothetic cues, from the environment (usually visual landmarks). In this paper we investigate how information from body-based idiothetic and visual allothetic cues are integrated for navigation.

Navigation using <u>only</u> body-based idiothetic cues (for example navigating in the dark) is called Path Integration. Path Integration is notoriously inaccurate involving both systematic and random errors [4–6]. For example, systematic error include biases induced by execution and past experiences such as history effects from past trials [7–10]. Random errors include noise in the body-based idiothetic sensory cues as well as in the integration process itself. These random errors accumulate with the square root of the distance and duration traveled in a manner similar to range effects in magnitude estimations; a consequence of the Weber–Fechner and Stevens’ Power Law [10–14]. Despite these sources of errors in path integration, humans and animals rely heavily on path integration because body-based idiothetic cues are constantly present (unlike visual allothetic landmark cues that may be sparse [6, 15]). In addition, path integration allows for flexible wayfinding by computing a route through new never experienced paths, and adjust for unexpected changes along the way [4, 16, 17].

Navigation using <u>only</u> visual allothetic cues (for example navigating a virtual world on a desktop computer) is called Map or Landmark Navigation [1, 18]. Pure landmark navigation (i.e. without body-based idiothetic cues) can only be studied in virtual environments, where body-based idiothetic cues can be decoupled from visual allothetic cues. In these studies, human participants show no differences in their navigational ability with or without isolation from body-based idiothetic cues, emphasizing that landmark navigation is a separate, and potentially independent computation from path integration [19].

Navigation using <u>both</u> body-based idiothetic and visual allothetic cues relies on both path integration and landmark navigation, yet exactly how the two processes work together is a matter of debate. In ‘cue combination’ (or ‘cue integration’) models, independent estimates from path integration and landmark navigation are combined to create an <u>average</u> estimate of location and heading. This averaging process is often assumed to be Bayesian, with each estimate weighed according to its reliability [20, 21]. Conversely, in ‘cue competition’ models, estimates from path integration and landmark navigation compete, with one estimate (often the more reliable) overriding the other completely. Based on this view, Cheng and colleagues proposed that path integration serves as a back-up navigational system that is used only when allothetic information is unreliable [22].

Empirical support exists for both cue combination and cue competition accounts. In a study by Chen and colleagues [23], humans in a virtual-navigation task averaged estimates from path integration and landmark navigation according to their reliability, consistent with a Bayesian cue combination strategy. Conversely, in a similar experiment by Zhao and Warren [24], participants primarily used visual allothetic information, often ignoring body-based idiothetic cues even when the mismatch was as large as 90°, consistent with a cue competition strategy. Similar discrepancies exist across the literature, with some studies supporting cue combination (and even optimal Bayesian cue combination) [23–27], and others more consistent with cue competition [24, 28–31].

Further complicating these mixed findings across studies are the large individual differences in navigation ability between participants [2, 32–34]. These individual differences encompass both high level processes, such as learning, knowledge, and decisions about routes [35–37], as well as lower level processes, such as how individuals respond to Corriolis forces and the perception of angular rotations due to differences in semi-circular canal radii [38, 39]. Such large individual differences also impact the integration of body-based idiothetic and visual allothetic cues and may be one reason for the discrepancies in the literature [23, 24].

In this paper, we investigate how people combine body-based idiothetic and visual allothetic cues in the special case of computing egocentric head direction. We focus on head direction because of its relative simplicity (compared to estimating both heading and location) and because the head direction system is known to integrate both vestibular (idiothetic) and visual (allothetic) cues [40]. In our task, participants performed full-body rotations to a goal with only a brief flash of visual feedback that either matched or mismatched their expectations. By building models of this task that capture the key features of cue combination and cue competition strategies, as well as the ‘pure’ strategies of path integration and landmark navigation, we find evidence for a hybrid strategy in which the estimates of path integration and landmark navigation are combined when the mismatch is small, but compete when the mismatch is large. Model comparison suggests that almost all participants use this strategy, with the large individual differences between participants being explained by quantitative differences in model parameters not qualitative differences in strategy. We therefore suggest that this flexible, hybrid strategy may underlie some of the mixed findings in the literature.

## Methods

### Participants

33 undergraduate students (18 female, 15 male, ages 18-21) received course credit for participating in the experiment. Of the 33, 3 students (3 female) did not finish block 1 due to cybersickness and were excluded from this study.

### Ethics statement

All participants gave written informed consent to participate in the study, which was approved by the Institutional Review Board at the University of Arizona.

### Stimuli

The task was created in Unity 2018.4.11f1 using the Landmarks 2.0 framework [41]. Participants wore an HTC Vive Pro with a wireless Adapter and held pair of HTC Vive controllers (Fig. 1A). The wireless headset, that was powered by a battery lasting about 2 hours, was tracked using 4 HTC Base Station 2.0, which track with an average positioning error of 17mm with 9 mm standard deviation [42]. Participants were placed in the center of a large rectangular (13m x 10m x 5m) naturalistic virtual room with several decorations (Fig. 1B). This was to ensure that visual feedback from different angles would be distinguishable by the geometry of the room and the decorations.

**Fig 1.**
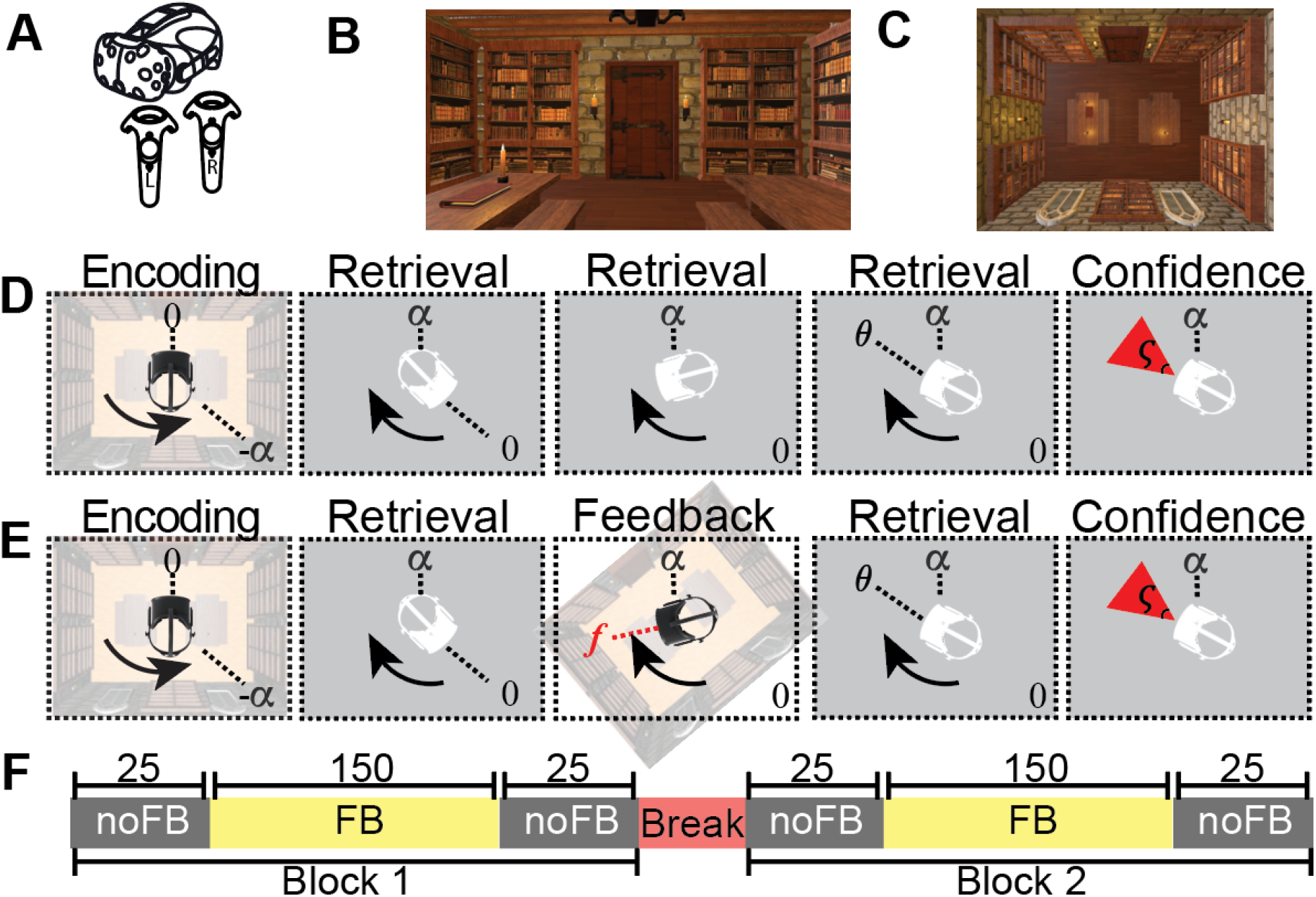
Task procedure. (A) Participants wear an HTC VIVE headset along with the handheld controllers to immerse themselves in a virtual room (B, C). First person view of the virtual environment at the beginning of a trial. (C) Top down view of the virtual environment. (D, E) Trial timeline for No Feedback (D) and Feedback (E) trials. At the start of each trial they face the door of the room and turn through −*α* degrees with visual feedback present. Visual feedback is then removed (gray squares) and they must turn back *α* degrees to face the door again. At the end of the turn participants stop at heading angle *θ*_*t*_ and report their confidence *ς* by adjusting the size of a red rectangle. The only difference between the No Feedback and Feedback conditions is the presence of a brief flash of visual information part way through the turn in the Feedback condition (E). Overall participants completed 100 trials of the Feedback condition and 300 trials of the No Feedback condition over the course of the experiment (F).

### Preprocessing and exclusion criteria

The 30 participants included for data analysis all completed block 1 (Fig. 1F). Due to tracking failures in the headset caused by a low battery, data from 3 participants was lost for most of session 2. Nevertheless we include data from all 30 participants in our analysis. Trials were removed if participants rotated in the wrong direction or if they pressed in the incorrect button to register their response. We also removed trials in which participants responded before they received feedback.

### The Rotation Task

During the procedure, participants were first guided through a rotation of −*α* degrees with visual feedback present (encoding phase). They were then asked to turn back to their initial heading, i.e. to turn a ‘target angle’ +*α*, with visual feedback either absent or limited (retrieval phase). In the No Feedback condition, participants received no visual feedback during the retrieval phase. In the Feedback condition, participants received only a brief flash of (possibly misleading) visual feedback at time *t*_*f*_. By quantifying the extent to which the feedback changed the participants’ response, the Rotation Task allowed us to measure how path integration is combining with visual feedback to compute heading.

More precisely, at the start of each trial, participants faced the door in a virtual reality room (Fig. 1B, C) and were cued to turn in the direction of the haptic feedback provided by a controller held in each hand (Fig. 1A). Feedback from the controller held in the left hand cued leftward rotations (counterclockwise) while feedback from the right controlled cued participants to rotate rightward (clockwise). Participants rotated until the vibration stopped at the encoding angle, −*α*, which was unique for each trial (sampled from a uniform distribution, 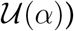). Participants were free to turn at their own pace and the experimenter provided no guidance or feedback on their rotational velocity. During the encoding procedure, participants saw the virtual room and were able to integrate both visual and vestibular information to compute their heading.

During the retrieval phase, participants had to try to return to their original heading direction (i.e. facing the door) with no (No Feedback condition) or limited (Feedback condition) visual feedback. At the beginning of the retrieval phase, participants viewed a blank screen (grey background in Fig. 1D). They then attempted to turn the return angle, +*α*, as best as they could based on their memory of the rotation formed during encoding. Participants received no haptic feedback during this retrieval process.

The key manipulation in this study was whether visual feedback was presented during the retrieval turn or not and, when it was presented, the extent to which the visual feedback was informative. In the No Feedback condition, there was no visual feedback and participants only viewed the blank screen — that is they could only rely on path integration to execute the correct turn. In the Feedback condition, participants saw a quick (300ms) visual glimpse of the room, at time *t*_*f*_ and angle *f*, which was either consistent or inconsistent their current bearing 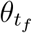. Consistent feedback occurred with probability *ρ* = 70%. In this case the feedback angle was sampled from a Gaussian centered at the true heading angle and with a standard deviation of 30°. Inconsistent feedback occurred with probability 1 − *ρ* = 30%. In this case the feedback was sampled from a uniform distribution between −180° and +180°. Written mathematically, the feedback angle, *f* was sampled according to

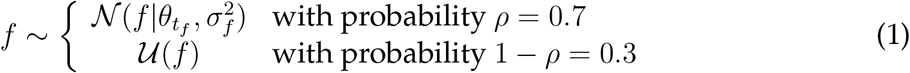

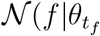 is a Gaussian distribution over *f* with mean 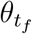 and standard deviation *σ*_*f*_ = 30°.

This form for the feedback sets up a situation in which the feedback is informative enough that participants should pay attention to it, but varied enough to probe the impact of misleading visual information across the entire angle space. To further encourage participants to use the feedback, they were not told that the feedback could be misleading.

Upon completing the retrieval turn, participants indicated their response with a button press on the handheld controllers (Fig. 1D), thus logging their response angle, *θ*_*t*_. Next, a red triangle appeared with the tip centered above their head and the base 6 meters away. Participants then adjusted the angle *ς* to indicate their confidence in their response angle using the touch pad on the controllers. In particular, they were told to adjust *ς* such that they were confident that the true angle *α* would fall within the red triangle (Fig. 1D). Participants were told they would received virtual points during this portion, with points scaled inversely by the size of the *ς* such that a small *ς* would yield to higher points (risky) and large *ς* would yield to lower points (safe).

After completing their confidence rating, the trial ended and a new trial began immediately. To ensure that participants did not receive feedback about the accuracy of their last response, each trial always began with them facing the door. This lack of feedback at the end of the trial ensured that participants were unable to learn from one trial to the next how accurate their rotations had been.

Overall the experiment lasted about 90 minutes. This included 10 practice trials (6 with feedback, 4 without) and 400 experimental trials split across two blocks with a 2-10 minute break between them (Fig. 1E). Of the 200 trials in each block, the first and last 25 trials in each block were No Feedback trials, while the remaining 150 were feedback trials. Thus each participant completed 100 trials in the No Feedback condition and 300 trials in the Feedback condition.

Participants were allowed to take a break at any time during the task by sitting on a chair provided by the experimenter. During these breaks participant continued to wear the VR headset and the virtual environment stayed in the same orientation.

## Models

We built four models of the Rotation Task that, based on their parameters, can capture several different strategies for integrating visual allothetic and body-based idiothetic estimates of location. Here we give an overview of the properties of these models, full mathematical details are given in the Supplementary Material.

### Path Integration model

In the Path Integration model we assume that the visual feedback is either absent (as in the No Feedback condition) or ignored (as potentially in some participants). In this case, the estimate of heading is based entirely on path integration of body-based idiothetic cues. To make a response, i.e. to decide when to stop turning, we assume that participants compare their heading angle estimate, computed by path integration, with their memory of the target angle. Thus, the Path Integration model can be thought of as comprising two processes: a path integration process and a target comparison process (Fig. 2).

**Fig 2.**
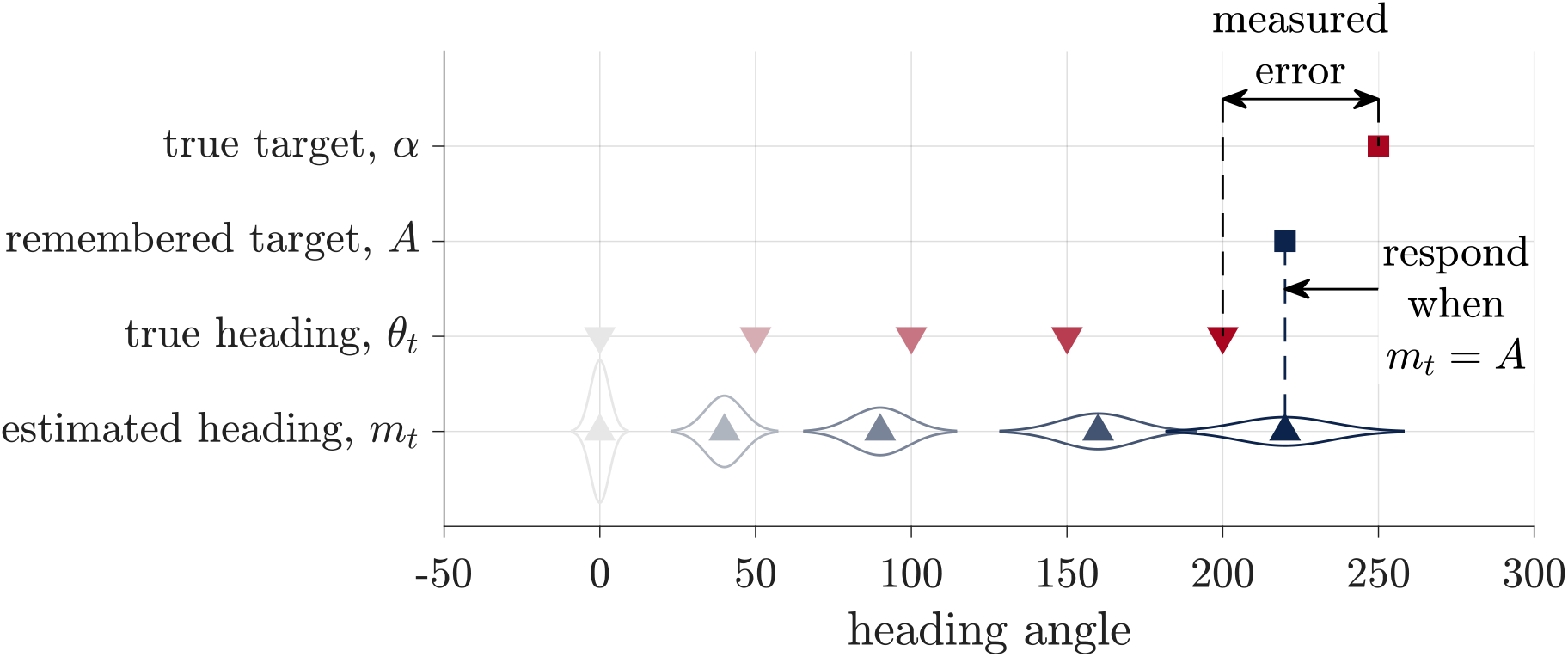
Schematic of the Path Integration model. During path integration, participants keep track of a probability distribution over their heading, which is centered at mean *m*_*t*_. To respond they compare this estimated heading to their remembered target location, *A*, halting their turn when *m*_*t*_ = *A*. The experimenter observers neither of these variables, instead we quantify the measured error as the difference between the true target angle, *α*, and the true heading angle, *θ*_*t*_.

In the path integration process, we assume that participants integrate biased and noisy vestibular cues about their angular velocity, *d*_*t*_. These noisy velocity cues relate to their true angular velocity, *δ*_*t*_ by

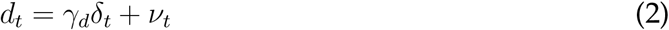

where *γ*_*d*_ denotes the gain on the velocity signal, which contributes to systematic under- or over-estimation of angular velocity and *ν*_*t*_ is zero-mean Gaussian noise with variance that increases in proportion to the magnitude of the angular velocity, |*δ*_*t*_|, representing a kind of Weber–Fechner law behavior [10].

We further assume that participants integrate this biased and noisy velocity information over time to compute a probability distribution over their heading. For simplicity we assume this distribution is Gaussian such that

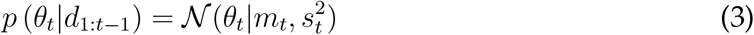

where the *m*_*t*_ is the mean of the Gaussian over heading direction and 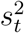 is the variance. Full expressions for *m*_*t*_ and 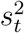 are given in the Supplementary Material. Fig. 2 illustrates how this distribution evolves over time.

In the target comparison process, we assume that participants compare their estimate of heading from the path integration process to their memory of the target angle. As with the encoding of velocity, we assume that this memory encoding is a noisy and biased process such that the participant’s memory of the target angle is

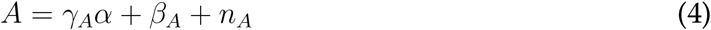

where *γ*_*A*_ and *β*_*A*_ are the gain and bias on the memory that leads to systematic over- or under-estimation of the target angle, and *n*_*A*_ is zero mean Gaussian noise with variance 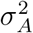.

To determine the response, we assume that participants stop moving when their current heading estimate matches the remembered angle. That is, when

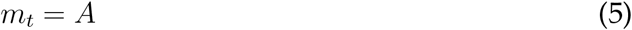

Substituting in the expressions for *m*_*t*_ and *A* (from the Supplementary Material), we can then compute the distribution over the measured error; i.e., the difference between participants actual heading and the target (*θ*_*t*_−*α*). Assuming all noises are Gaussian implies that the distribution over measured error is also Gaussian with a mean and variance given by

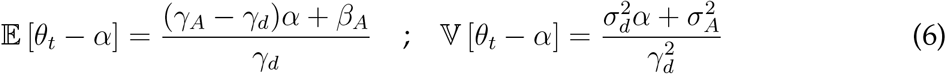

Thus, the Path Integration model predicts that both the mean error and the variance in the mean error will be linear in the target angle, *α*, a prediction that we can test in the No Feedback condition (Fig. 7). In addition, in the Feedback condition, the Path Integration model also predicts that the response error in the Feedback condition will be independent of the visual feedback (Fig. 6A), a result that should not be surprising given that the Path Integration model ignores visual feedback.

### Kalman Filter model

Unlike the Path Integration model, which always ignores feedback, the Kalman Filter model always incorporates the visual feedback into its estimate of heading (Fig. 3). Thus the Kalman Filter model captures one of the key features of the Landmark Navigation strategy. However, it is important to note that the Kalman Filter model is slightly more general than ‘pure’ Landmark Navigation. Indeed, for most parameter values, it is a cue combination model in that it combines the the visual feedback with the estimate from Path Integration. Only for some parameter settings (as we shall see below), does the Kalman Filter model converge to a pure Landmark Navigation strategy in which it completely ignores prior idiothetic cues when visual feedback is presented.

**Fig 3.**
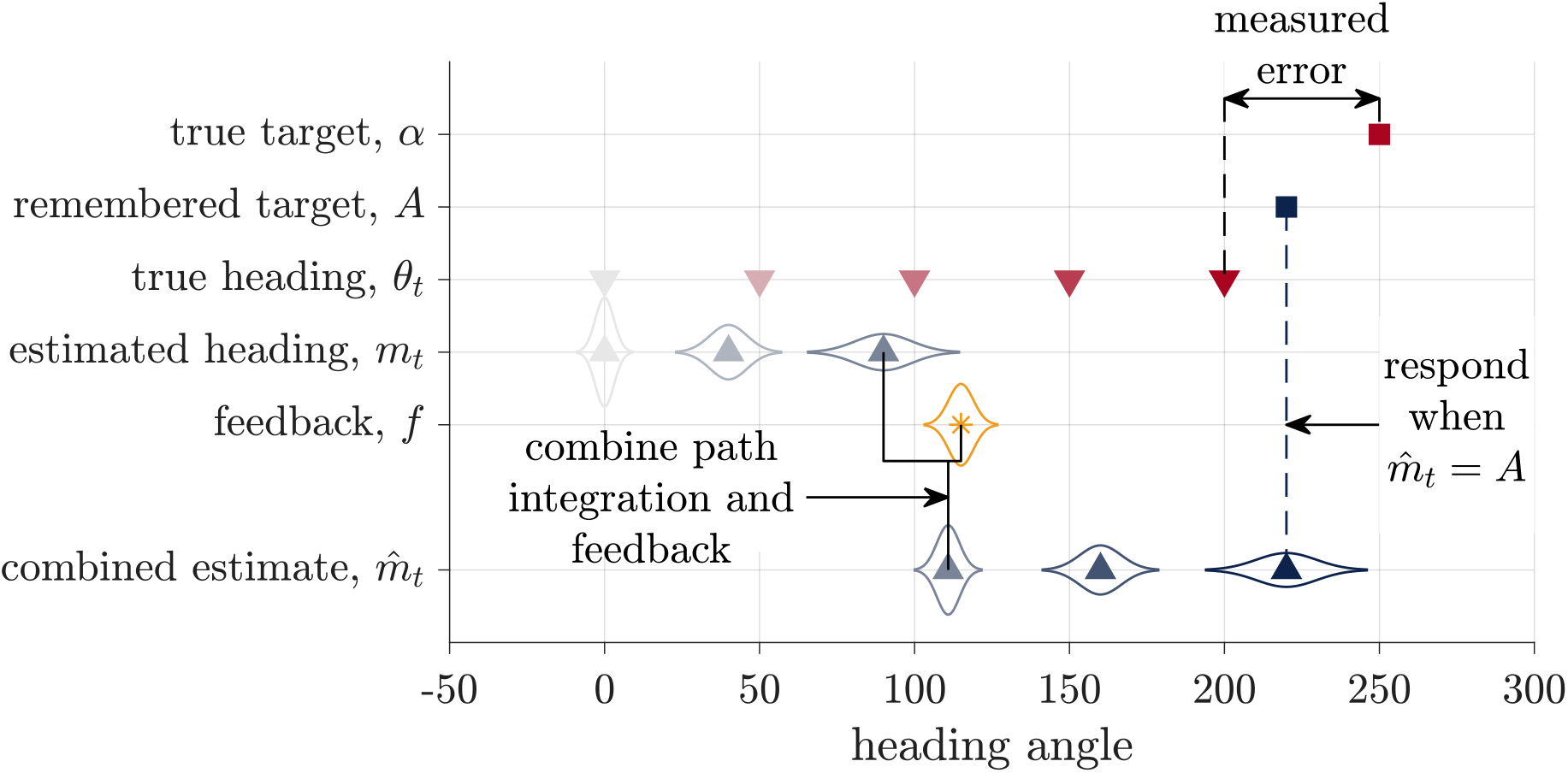
Schematic of the Kalman Filter model. Similar to the Path Integration model, this models assumes that participants keep track of a probability distribution over their heading that, before the feedback, is centered on mean *m*_*t*_. When the feedback, *f*, is presented, they combine this visual information with their path integration estimate to compute a combined estimate of heading 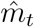. They then stop turning and register their response when 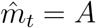, their remembered target. As with the Path Integration model, none of these internal variables are observed by the experimenter, who instead measures the error as the difference between the true target, *α*, and heading angle *θ*_*t*_.

The Kalman Filter model breaks down the retrieval phase of the task into four different stages: initial path integration, before the visual feedback is presented; feedback incorporation, when the feedback is presented; additional path integration, after the feedback is presented; and target comparison, to determine when to stop.

Initial path integration, is identical to the Path Integration model. The model integrates noisy angular velocity information over time to form an estimate of the mean, *m*_*t*_ and uncertainty, *s*_*t*_, over the current heading angle *θ*_*t*_.

When feedback (*f*) is presented, the Kalman Filter model incorporates this feedback with the estimate from the initial path integration process in a Bayesian manner. Assuming all distributions are Gaussian, the Kalman Filter model computes the posterior distribution over head direction as

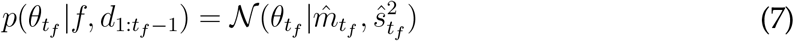

where 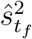 is the variance of the posterior (whose expression is given in the Supplementary Material) and 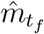 is the mean given by

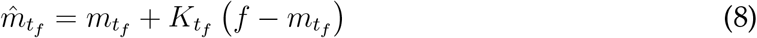

where 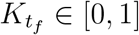 is the ‘Kalman gain,’ sometimes also called the learning rate [43, 44].

The Kalman gain is a critical variable in the Kalman Filter model because it captures the relative weighting of idiothetic (i.e. the estimate from Path Integration, 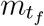) and allothetic (i.e. the feedback, *f*) information in the estimate of heading. In general, the Kalman gain varies from person to person and from trial to trial depending on how reliable people believe the feedback to be relative to how reliable they believe their path integration estimate to be. When the model believes that the Path Integration estimate is more reliable, the Kalman gain is closer to 0 and idiothetic cues are more heavily weighted. When the model believes that the feedback is more reliable, the Kalman gain is closer to 1 and the allothetic feedback is more heavily weighted. In the extreme case that the model believes that the feedback is perfect, the Kalman gain is 1 and the Kalman Filter model implements ‘pure’ landmark navigation, basing its estimate of heading entirely on the visual feedback and ignoring the path integration estimate completely.

After the feedback has been incorporated, the model continues path integration using noisy velocity information until its estimate of heading matches the remembered target angle. Working through the algebra (see Supplementary Material) reveals that the measured response distribution is Gaussian with a mean given by

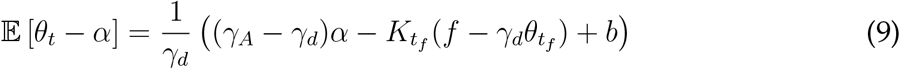

This implies that the error in the Kalman Filter model is linear in the feedback prediction error 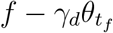 (Fig. 6B).

### Cue Combination model

Like the Kalman Filter model, the Cue Combination model combines the feedback with the estimate of heading from path integration. Unlike the Kalman Filter model, however, the Cue Combination model also takes into account the possibility that the feedback will be misleading, in which case the influence of the feedback is reduced. In particular, the Cue Combination model computes a mixture distribution over heading angle with one component of the mixture assuming that the feedback is false and the other that the feedback is true. These two components are weighed according to the computed probability that the feedback is true, *p*_*true*_ (Fig. 4).

**Fig 4.**
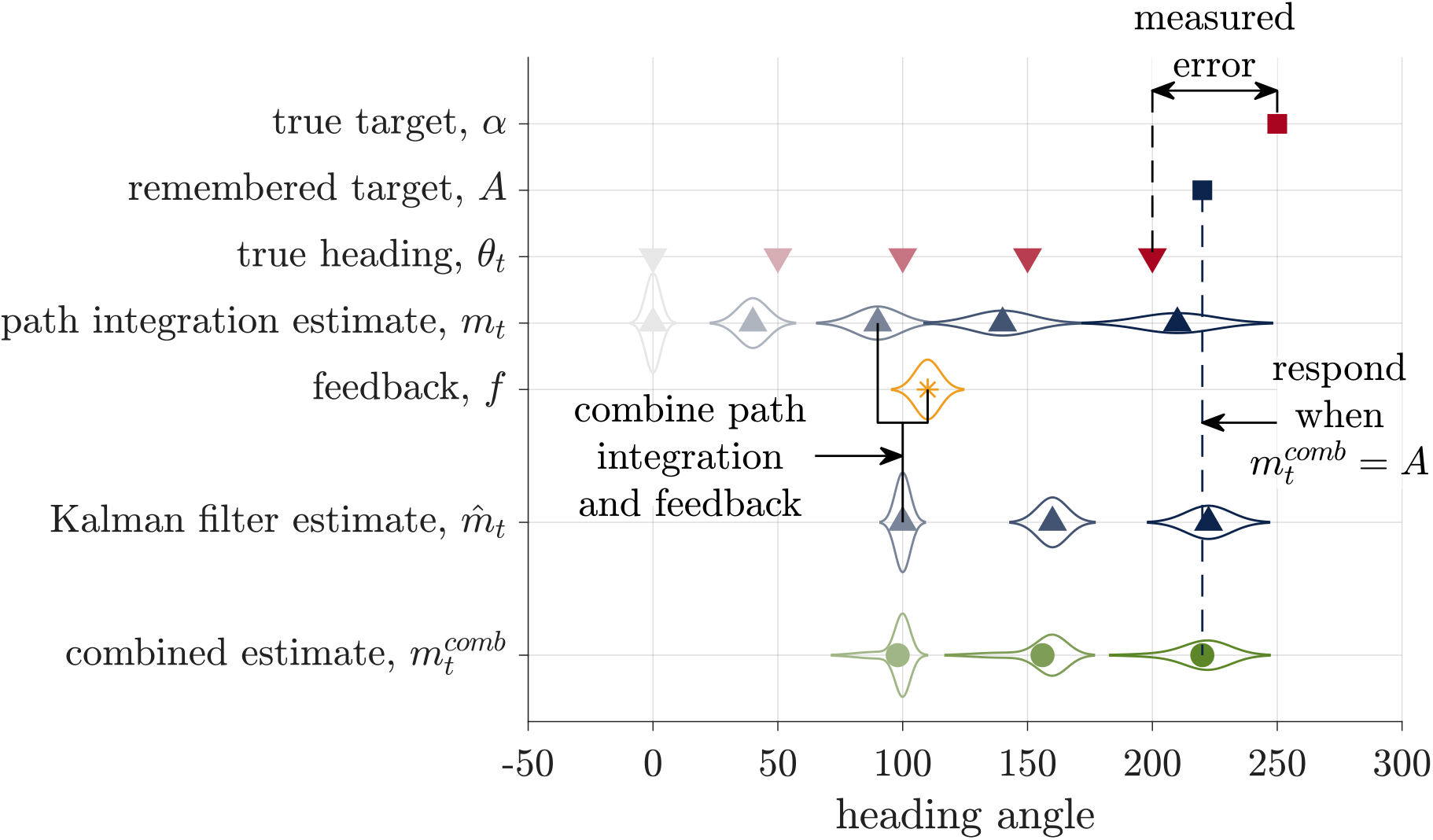
Schematic of the Cue Combination model. The Cue Combination model combines two estimates of heading, the path integration estimate, *m*_*t*_, and the Kalman Filter estimate, 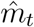, to compute a combined estimate 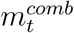. The response is made when this combined estimate matches the remembered target *A*.

Mathematically, the Cue Combination model computes the probability distribution over heading angle by marginalizing over the truth of the feedback

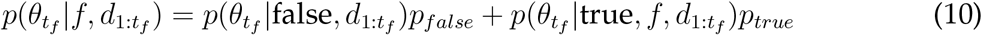

where 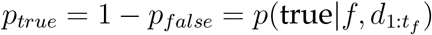 is the probability that the feedback is true given the noisy velocity cues seen so far. Consistent with intuition, *p*_*true*_ decreases with the absolute value of the prediction error at the time of feedback 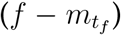 such that large prediction errors are deemed unlikely to come from true feedback.

Equation 10 implies that, at the time of feedback, the Cue Combination model updates its estimate of the mean heading by <u>combining</u> the estimates from Path Integration model, 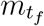, with the estimate from the Kalman Filter model, 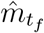, as

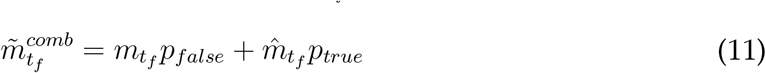

Assuming that a similar target comparison process determines the response (i.e. participants stop turning when 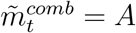), then this implies that the response distribution for the Cue Combination model will be Gaussian with a mean error given by the mixture of the Path Integration and Kalman Filter responses:

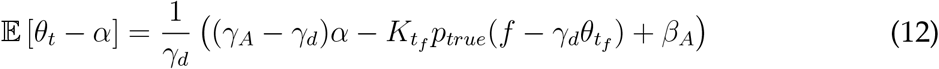

Because *p*_*true*_ depends on the prediction error, Equation 12 implies that the average error in the Cue Combination has a non-linear dependence on the prediction error (Fig. 6C).

### Hybrid model

Instead of averaging over the possibility that the feedback is true or false, in the Hybrid model the estimates from the Path Integration model and the Kalman Filter model compete (Fig. 5). This is because the Kalman Filter model is itself a cue combination model the Hybrid model is a true hybrid between cue combination and cue competition.

**Fig 5.**
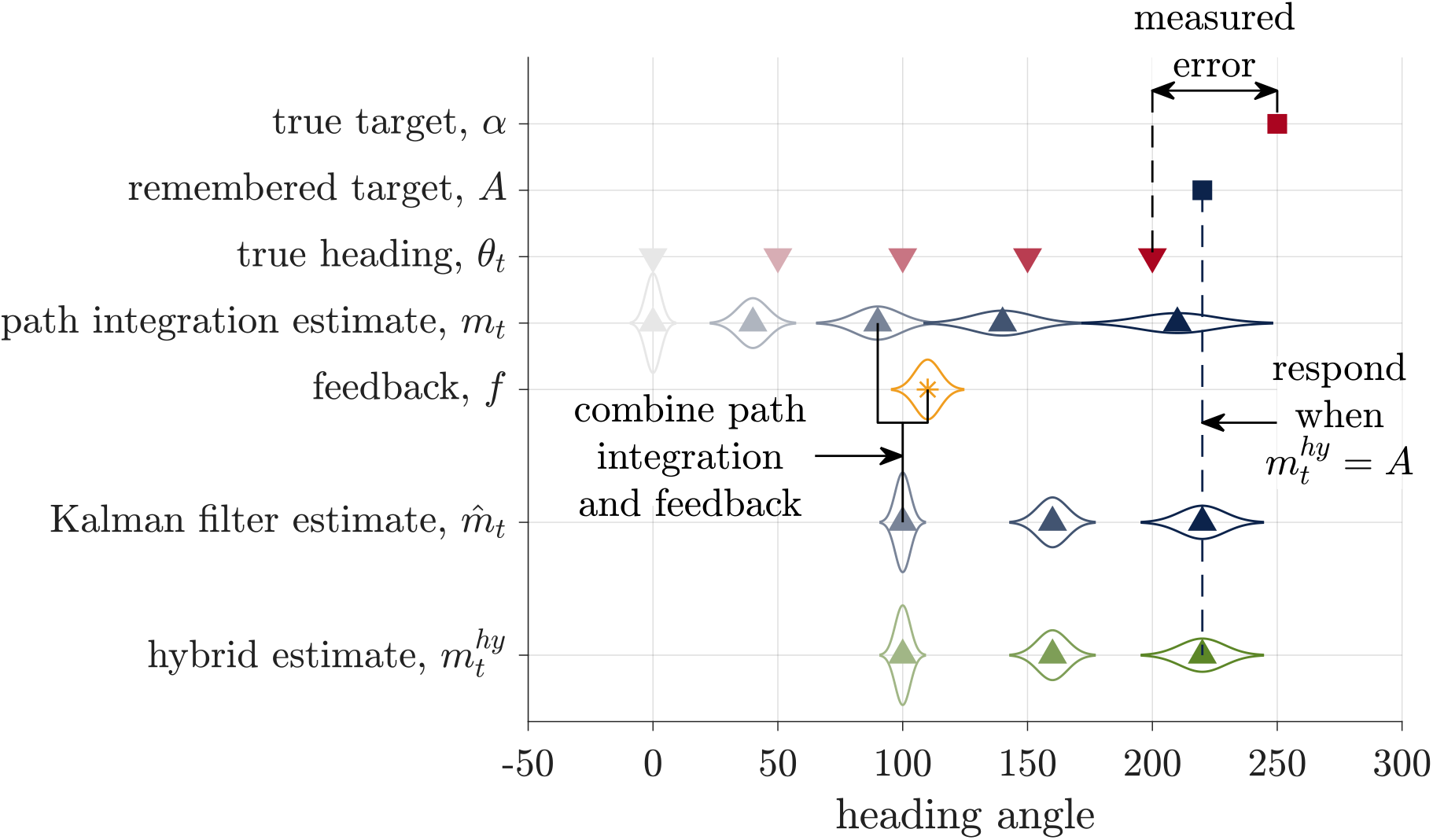
Schematic of the Hybrid model. The Hybrid model bases its estimate of heading, 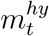, either on the path integration estimate or the Kalman Filter estimate. Here we illustrate the case where the model chooses the landmark navigation estimate. The response is made when the hybrid estimate matches the remembered target angle.

In particular, we assume that the Hybrid model makes the decision between Path Integration and Kalman Filter estimates according to the probability that the feedback is true (*p*_*true*_), by <u>sampling</u> from the distribution over the veracity of the feedback. Thus with probability *p*_*true*_, this model behaves exactly like the Kalman Filter model, setting its estimate of heading to 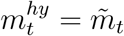, and with probability *p*_*false*_ = 1 − *p*_*true*_ this model behaves exactly like the Path Integration model, setting its estimate of heading to 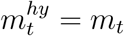. This implies that the distribution of errors is a mixture of the Kalman Filter and Path Integration models such that the average response error is

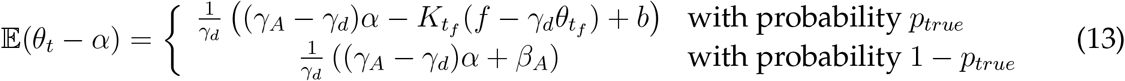

This competition process ensures that the relationship between response error and feedback offset will be a mixture of the Path Integration and Kalman Filter responses (Fig. 6D). When the model decides to ignore the feedback, the response will match the Path Integration model. This occurs most often for large offset angles, when *p*_*true*_ is closer to 0. When the model decides to incorporate the feedback, the response will lie on the red line. This occurs most often for small offset angles, when *p*_*true*_ is closer to 1.

**Fig 6.**
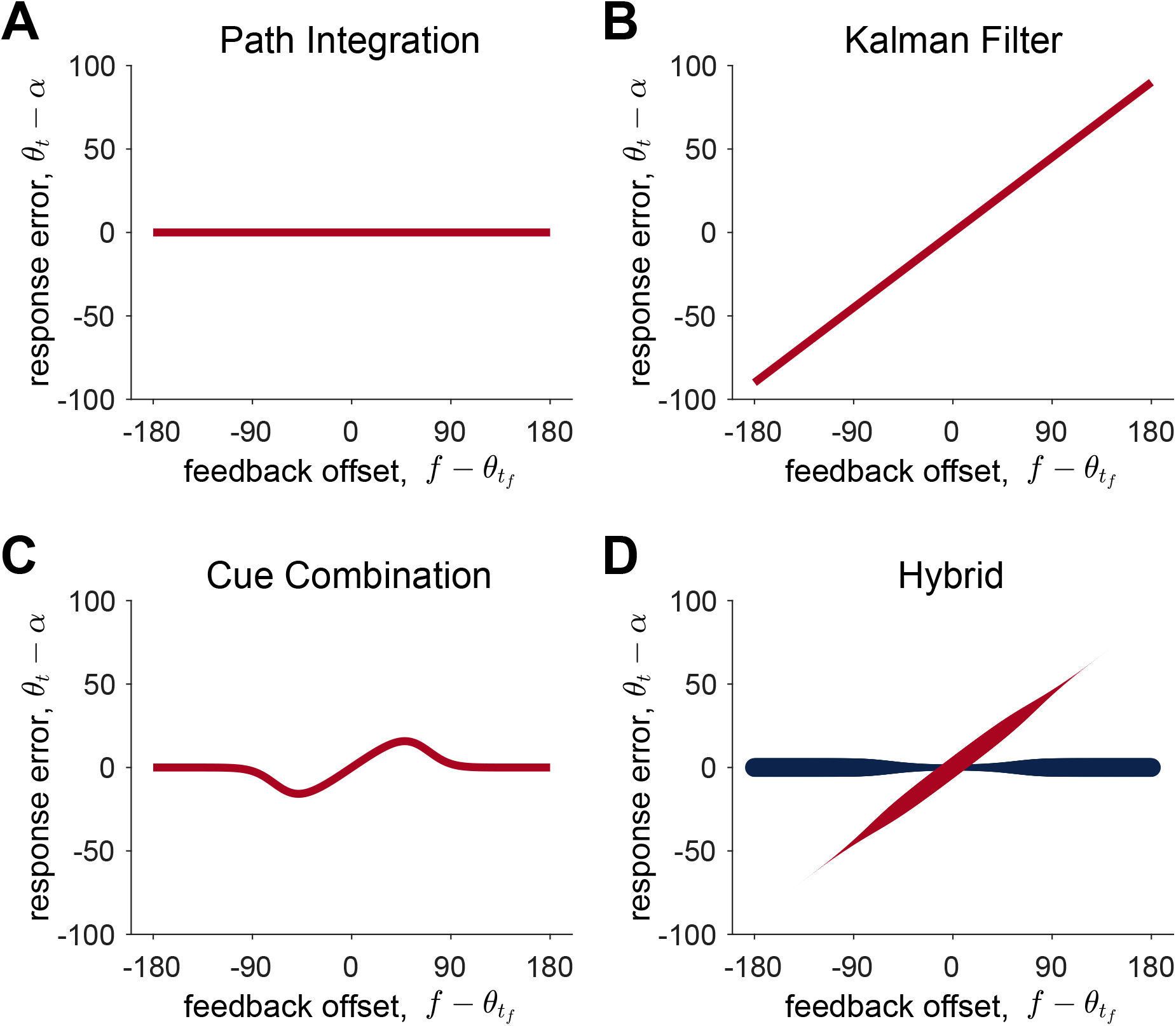
Model predictions for the Path Integration, Kalman Filter, Cue Combination, and Hybrid models. In (A-C) the red lines correspond to the mean of the response error predicted by the model. In (D) the two lines correspond to the mean response when the model assumes the feedback is true (red) and false (blue). The thickness of the red and blue lines in (D) corresponds to the probability that the model samples from a distribution with this mean, i.e. *p*_*true*_ for red and 1 − *p*_*true*_ = *p*_*false*_ for blue.

### Model fitting and comparison

Each model provides a closed form function for the likelihood that the a particular angular error is observed on each trial, *τ*, given the target, the feedback (in the Feedback condition), and the true heading angle at feedback. That is, we can formally write the likelihood of observing error^*τ*^ on trial *τ* as

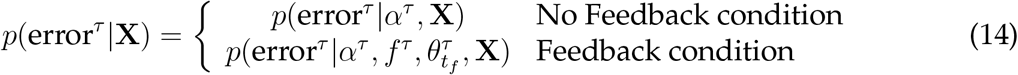

where vector **X** denotes the free parameters of the model. In all cases we used the Path Integration model to compute the likelihoods on the No Feedback trials and each of the four models to compute the likelihoods on the feedback trials. When combining each model with the Path Integration model in this way, we yoked the shared parameters between the models to be equal across the No Feedback and Feedback trials.

We then combined the likelihoods across trials to form the log likelihood for a given set of parameters

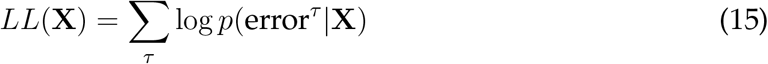

where the sum is over the trials in both the No Feedback and Feedback conditions. The best fitting parameters were then be computed as those that maximize this log likelihood

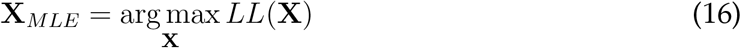

Model fitting was performed using the fmincon function in Matlab. To reduce the possibility of this optimization procedure getting trapped in local minima, we ran this process 100 times using random starting points. Each starting point was randomly sampled between the upper and lower bound on each parameter value as defined in Table S1. Parameter recovery with simulated data showed that this procedure was able to recover parameters adequately for all models (Supplementary Section 3, and Figs. S6, S8, S9, and S10).

Model comparison was performed by computing the Bayes Information Criterion (BIC) for each model for each participant

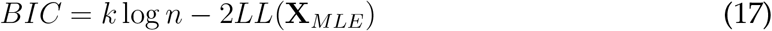

where *k* is the number of free parameters in the model and *n* is the number of trials in the data. Model recovery with simulated data showed that this procedure was sufficient to distinguish between the four models on this experiment (Supplementary Section 3 and Fig. S4).

## Results

### Behavior in the No Feedback condition is consistent with the Path Integration model

The Path Integration model predicts that the mean of the response error will be linear in the target angle *α*. To test whether this linear relationship holds, we plotted the response error *θ*_*t*_ − *A* as a function of target angle *α* for all of the No Feedback trials (Fig. 7). This reveals a clear linear relationship between the mean response error and target angle. In addition, for many participants, the variability of the response error also appears to increase with the target angle, which is also consistent with the Path Integration model. Notable in Fig. 7 are the considerable individual differences between participants, with some participants having a negative slope (systematically underestimating large target angles), some a positive slope (overestimating large target angles), and some with approximately zero slope.

**Fig 7.**
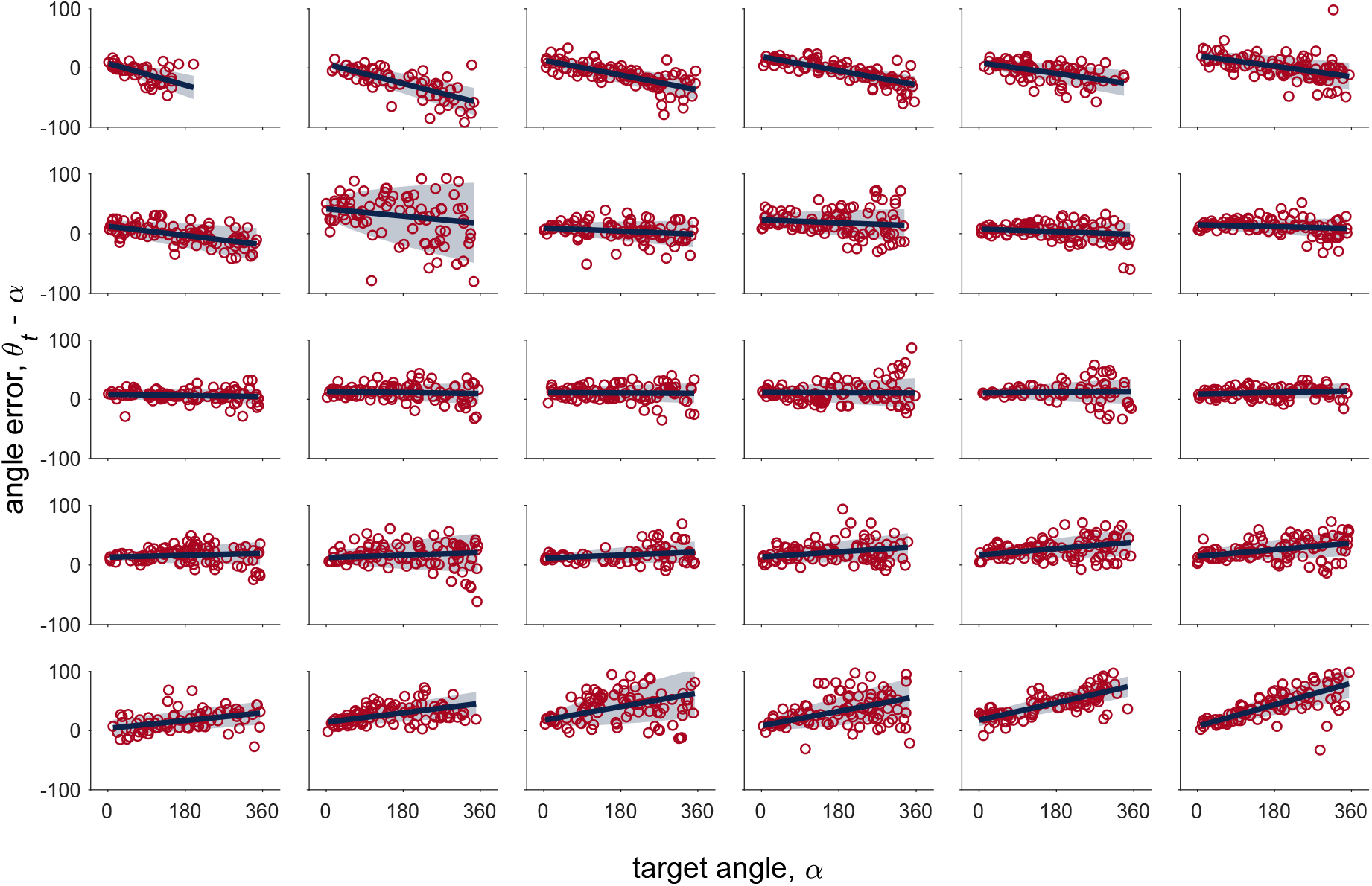
Error vs target angle for the No Feedback condition. Each plot corresponds to data from one participant and plots are ordered from most negative slope (top left) to most positive slope (bottom right). The red circles correspond to human data, the solid blue to the mean error from the Path Integration model fit, and shaded blue area to the mean ± standard deviation of the error from the Path Integration model fit.

To investigate further, we fit the Path Integration model to the No Feedback data. This model has five free parameters capturing the gain and noise in the velocity signal (*γ*_*d*_, *σ*_*d*_) and the gain, bias and noise in the target encoding process (*γ*_*A*_, *β*_*A*_, *σ*_*A*_). Because *γ*_*d*_ only appears as part of a ratio with other parameters in the Path Integration model, it cannot be estimated separately. We therefore fix the value of the velocity gain to *γ*_*d*_ = 1 and interpret the resulting parameter values as ratios (e.g. *γ*_*A*_/*γ*_*d*_ etc …).

As shown in Fig. 7, the Path Integration model provides an excellent fit to the No Feedback data, accounting for both the linear relationship between response error and target and the increase in variability with target.

Looking at the best fitting parameter values, we find that the target gain is close to 1 at the group level (mean *γ*_*A*_/*γ*_*d*_ = 0.997), indicating no systematic over- or under-weighting of the target across the population (Fig. 8A). Individual participants vary considerably, however, with *γ*_*A*_/*γ*_*d*_ ranging from 0.8 (negative slope in Fig. 7) to 1.2 (positive slope in Fig. 7). In contrast to the target gain, we find a systematic target bias across the population, with all participants turning slightly too far (mean *β*_*A*_/*γ*_*d*_ = 13.4°; Fig. 8B). Nonetheless, as can be seen in 7, there is considerable variability across participants.

**Fig 8.**
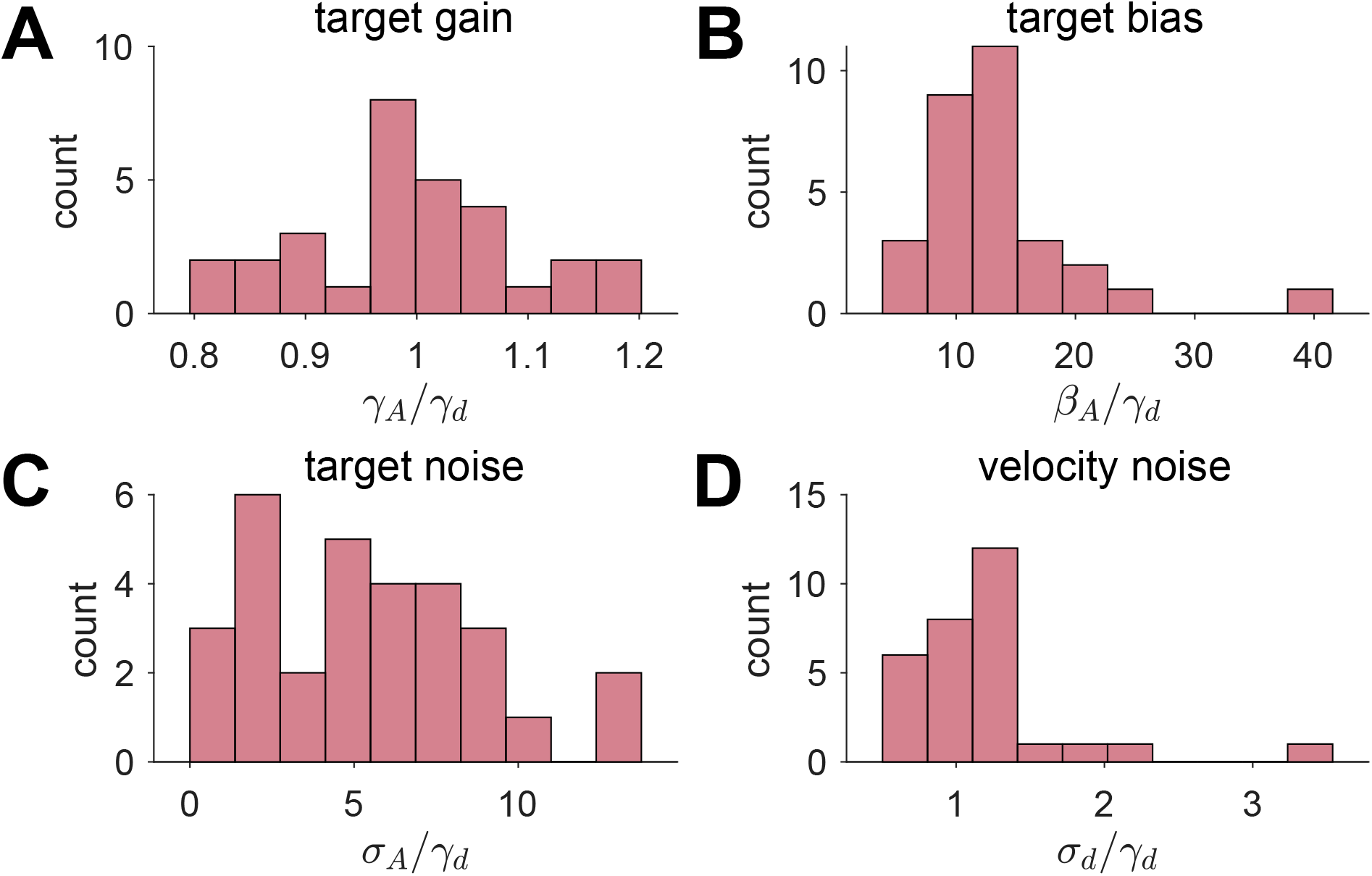
Parameter values for the Path Integration model fit to the No Feedback data.

For accuracy, we find that most participants have some target noise (mean *σ*_*A*_/*γ*_*d*_ = 5.35°; Fig. 8C) and all participants have velocity noise (mean *σ*_*d*_/*γ*_*d*_ = 1.19°; Fig. 8D). This latter result suggests that the variance of the noise in head direction estimates grows linearly with target angle and with a constant of proportionality close to 1.

### Behavior in the Feedback condition is consistent with the Hybrid model

The key analysis for the Feedback condition relates the feedback offset, 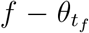, to the response error, *θ*_*t*_ − *α*. As illustrated in Fig. 6, each model predicts a different relationship between these variables.

In our experiment we found examples of behavior that was qualitatively consistent with all four models. These are illustrated in Fig. 9. At the extremes, Participant 27 appeared to ignore feedback completely, like what is shown for the path integration model (Fig. 9A), while Participant 30 seemed to always use the feedback, just like the Kalman Filter model (Fig. 9B). Conversely, Participant 15 appeared to use a Cue Combination approach, while Participant 2’s behavior was more consistent with the Hybrid model. This latter behavior is especially interesting because it strongly suggests a bimodal response distribution for large feedback offsets.

**Fig 9.**
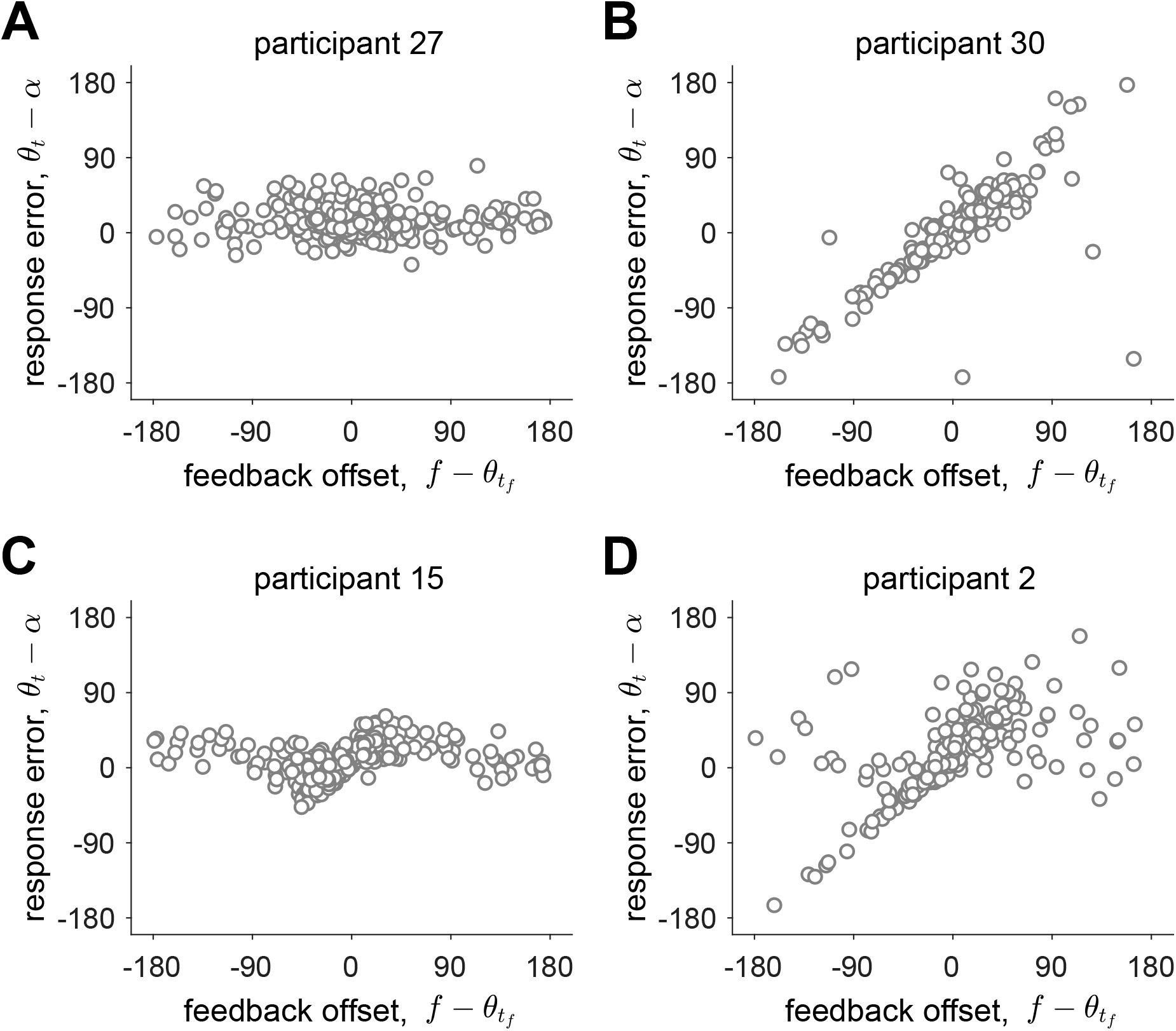
Examples of human behavior on the feedback trials.

To quantitatively determine which of the four models best described each participant’s behavior we turned to model fitting and model comparison. We computed Bayes Information Criterion (BIC) scores for each of the models for each of the participants and asked which model had the lowest BIC score for each person. Fig. 10A plots BIC scores for each model relative to the BIC score for the Hybrid model for each participant. In this plot, positive values correspond to evidence in favor of the Hybrid model, negative values correspond to evidence in favor of the other models. As can be seen in 10, the the Hybrid model is heavily favored and best describes the behavior of all but three participants (participant 27, who is best fit by the Path Integration model, and participants 15 and 23, who are best fit by the Cue Combination model; Fig. 10B).

**Fig 10.**
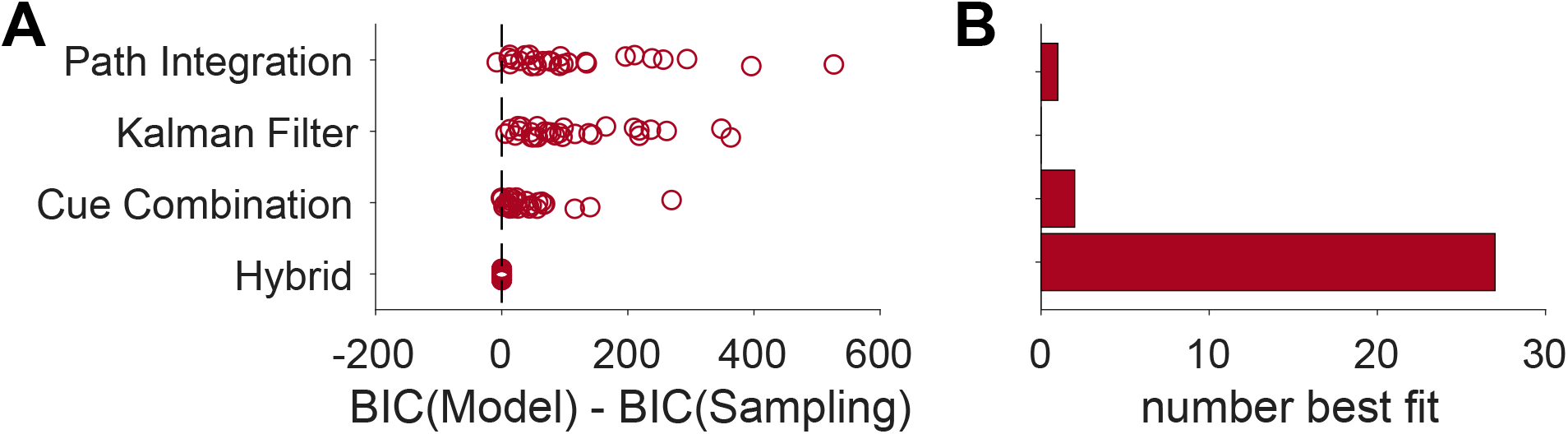
Model comparison. (A) BIC scores for each model relative to the BIC score for the Hybrid model for each participant. For each model, each circle corresponds to one participant. Positive numbers imply the fit favors the Hybrid model, negative numbers imply that the fit favors the other model. (B) The number of participants best fit by each model. 28 out of 30 participants were best fit by the Hybrid model, suggesting that this model best describes human behavior.

Qualitatively, the Hybrid model provides a good account of the data despite the large individual differences in behavior. In Fig. 11 we compare the behavior of the model to the behavior of four example participants. As already suggested in Fig. 9, Participant 2 is one of the cleanest examples of Hybrid behavior and it is not surprising that this behavior is well described by the model. Likewise the Hybrid model does an excellent job capturing the behavior of Participant 30, whose qualitative behavior appears more Kalman Filter like. The reason the Hybrid model outperforms the Kalman Filter model for this participant is that Participant 30 appears to ignore the stimulus on two trials at offsets of around −100 and +100 degrees. These data points correspond to large deviations from the Kalman Filter model behavior but are a natural consequence of the Hybrid model. The Hybrid model also captures the behavior of participants who integrate the feedback over a much smaller range such as Participants 10 and 25. A comparison between the Hybrid model and all participants is shown in Supplementary Fig. S13.

**Fig 11.**
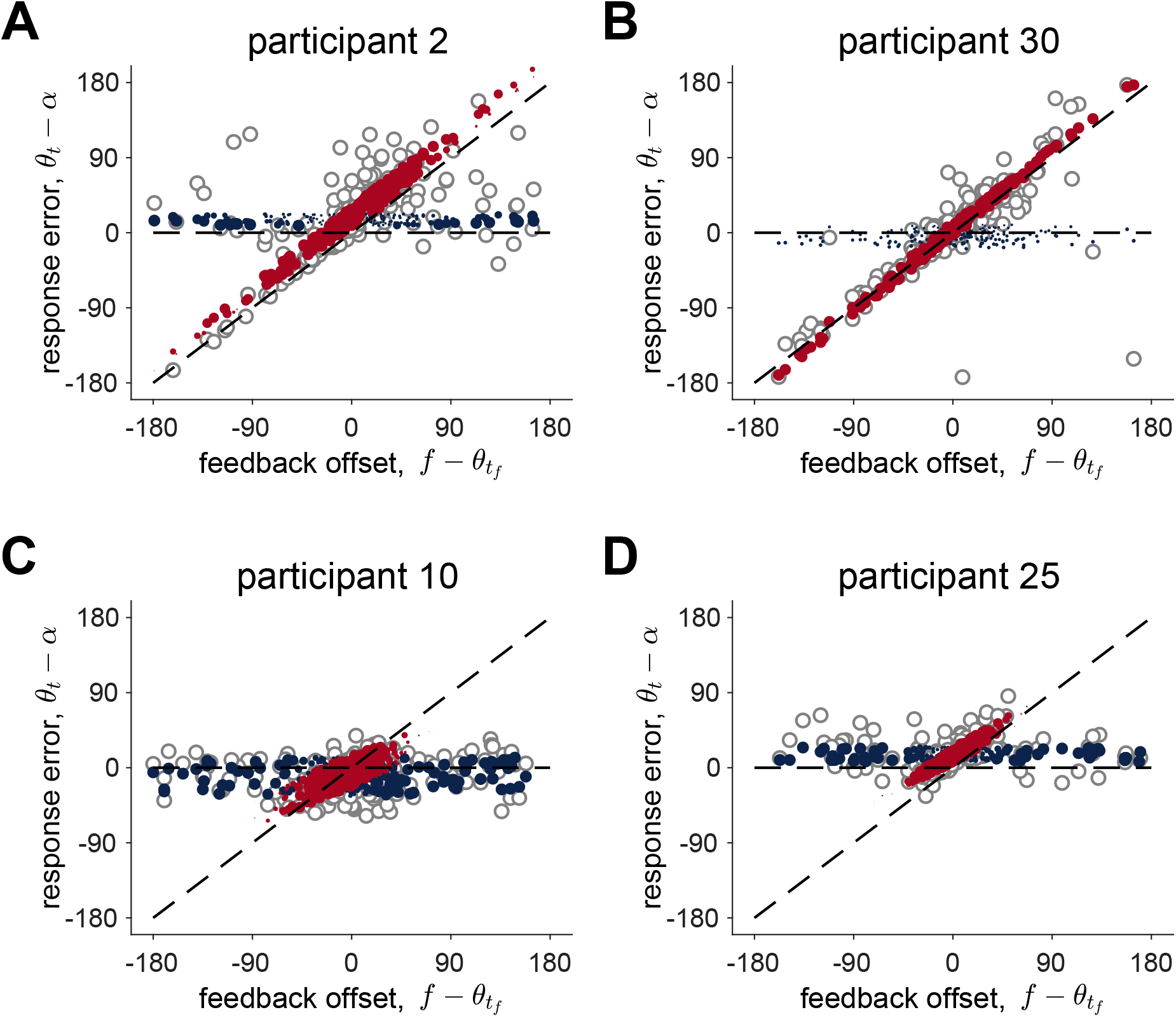
Comparison between data and the Hybrid model for four participants.

### Parameters of the Hybrid model suggest people use a true hybrid strategy between cue combination and cue competition

Consistent with the individual differences in behavior, there were significant individual differences in the fit parameter values across the group (Supplementary Fig. S11). Of particular interest is what these parameter values imply for the values of the Kalman gain, 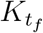. As mentioned in the Methods section, this variable is important because it determines the extent to which the Kalman Filter component of the Hybrid model incorporates the allothetic visual feedback vs the idiothetic path integration estimate of heading. The larger the Kalman gain, the more allothetic information is favored over idiothetic information. Moreover, if the Kalman gain is 1, then the Hybrid model becomes a ‘pure’ cue competition model. This is because the Kalman Filter component of the model ignores idiothetic information prior to the feedback (i.e. it implements ‘pure’ landmark navigation). Thus, the Hybrid model now decides between pure Path Integration and pure Landmark Navigation, consistent with a pure Cue Combination approach.

In Fig. 12, we plot implied Kalman gains for all trials for each participant. This clearly shows that the majority of participants do not have 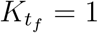, instead showing intermediate values for the Kalman gain. Thus we conclude that participants use a true hybrid of cue combination, when the mismatch between idiothetic and allothetic information is small, and cue competition when the mismatch is large.

**Fig 12.**
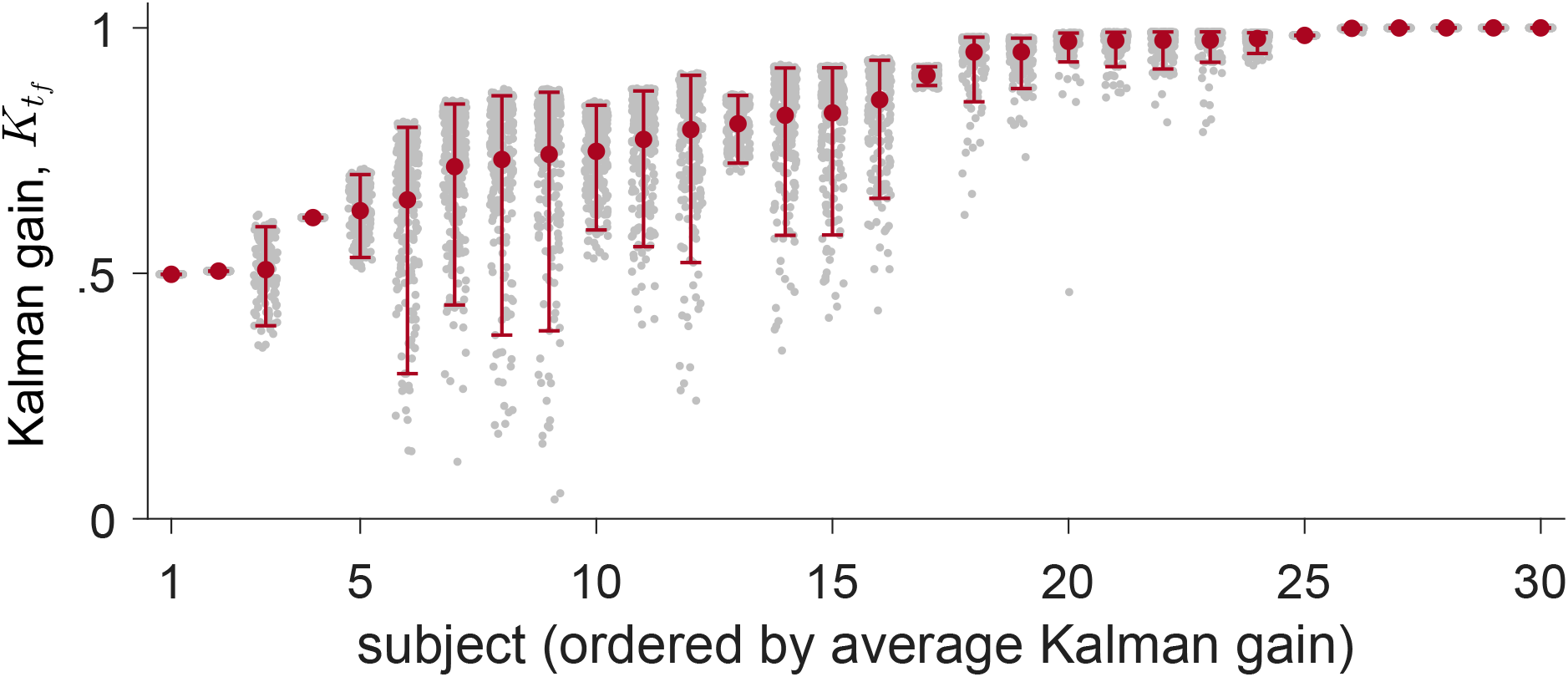
Computed Kalman gain for all participants and all trials. The Kalman gains computed for each trial for each participant are shown as gray dots. The mean Kalman gain and 95% confidence intervals are shown in red.

## Discussion

In this paper, we investigated how humans integrate path integration (rotational) and visual landmarks/ boundaries to estimate their heading. In our experiment, The Rotation Task, participants made a series ‘of turns in virtual reality, mostly without visual feedback. Visual feedback, when it was presented in the form of the boundaries of the room with distinct landmarks, was brief and offset from the true heading angle. This offset led to systematic errors in people’s turning behavior that allowed us to quantify how people combine visual allothetic feedback with their internal estimate of heading direction, computed by path integration of body-based idiothetic cues.

While there were considerable individual differences in task performance, our findings suggest that the majority of participants used the same overarching <u>hybrid</u> strategy to complete the task. In this strategy, body-based idiothetic and visual allothetic cues are combined when the estimates of path integration and landmark navigation are close and compete when the estimates are far apart. This behavior was well accounted for by a computational model that switches between competition and combination according to the subjective probability that the feedback is valid.

These findings support a hybrid model which may help to explain the mixed reports in the literature regarding cue combination. Specifically, some studies report evidence of cue combination [23, 25, 28, 45] while find evidence for others cue competition [24, 46]. One set of studies using a similar experimental paradigm involves the short-range homing task of Nardini and colleagues and shows evidence for both cue combination and competition depending on the conditions tested [25]. In this task, participants picked up three successive objects in a triangular path and returned them after a delay. During the return phase of the task, the experimenters manipulated the visual feedback to induce a 15 degree mismatch with the body-based cues. When both visual and body-based cues were present, Nardini et al. found that the variance of the response was smaller than when navigation relied on only one cue, consistent with the combination of visual and body-based cues in a Bayesian manner. However, when Zhao and Warren [24] increased the offset from 15 to 135 degrees, they found that participants based their estimate of location either entirely on the visual cues (when the offset was small) or entirely on the body-based cues (when the offset was large), taking this as evidence for a cue-competition strategy. Thus, in the same task, participants appeared to switch from cue combination to cue competition as the offset grew larger, exactly what we observe in our experiment, and what is predicted by the Hybrid model.

More generally, our model fits with bounded rationality theories of human cognition [47–50]. That is, people have limited computational resources, which in turn impacts the kinds of computations they can perform and how they perform them. In our case, combining visual and body-based cues to compute a probability distribution over heading should be easier when the cues align and the distribution is unimodal than when the cues conflict and the posterior is bimodal. In this latter case, representing the bimodal distribution by sampling one mode or the other, as the Hybrid model does, may be a rational strategy that demands fewer computational resources. Indeed, other work has shown that people may represent complex and multimodal distributions with a small number of samples, which may be as low as just a single sample in some cases [51–53].

A key prediction of such a sampling interpretation of the Hybrid model is that participants should sometimes lose information when integrating visual allothetic and body-based idiothetic cues. That is, when faced with a large feedback offset, instead of computing the full posterior distribution over heading, participants collapse this bimodal distribution to a unimodal distribution centered on the estimate from path integration or landmark navigation.

Such a ‘*semi*-Bayesian’ interpretation, stands in contrast to a fully-Bayesian alternative in which, participants do indeed keep track of the bimodal posterior and instead sample their estimate of heading direction from this posterior to determine their response. In this view, when faced with a large feedback offset, participants do compute the full distribution over heading, but rather than average over this distribution to compute their response, they sample from it instead. This implies that participants do not make a decision to ignore or incorporate the feedback and, as a result, do not lose information about the stimulus or their path integration estimate.

A key question for future work will be to distinguish between these two interpretations of the task. Does sampling occur at the time of feedback causing a collapse of the posterior distribution to one mode and a loss of information? Or does sampling occur later on and without the collapse of the posterior? Both interpretations lead to identical behavior on the Rotation Task. However, a modified version of the task should be able to distinguish them.

A key goal for future work will be to expand our experimental paradigm to test whether the hybrid model can generalize to navigational behavior from both rotational and translational movements. Another future gold will be to combine the Rotation Task with physiological measures (such as EEG) to study the neural underpinnings of this process [54]. In addition, it will be interesting to explore individual differences in behavior in more diverse populations including older adults and people with psychiatric disorders. By providing within-trial dynamics of cognitive variables as well as characterizing large individual differences with different parameter values, our task and model could help to set the stage for this future work.

## Supporting information

**S1 Fig. Graphical representation of the Path integration model**

**S2 Fig. Graphical representation of the Kalman Filter model.**

**S3 Fig. Graphical representation of the Cue Combination and Hybrid models.**

**S4 Fig. Model recovery confusion matrix.**

**S5 Fig. Parameter recovery for Path Integration model.**

**S6 Fig. Parameter recovery for Hybrid mode.**

**S7 Fig. No induced correlations.**

**S8 Fig. Parameter recovery for Path Integration model.**

**S9 Fig. Parameter recovery for Kalman Filter model.**

**S10 Fig. Parameter recovery for Cue Combination model.**

**S11 Fig. Best fit parameters for the Hybrid model.**

**S12 Fig. Correlations between parameters in the Hybrid model.**

**S13 Fig. Comparison between data and model for all participants.**

**S14 Fig. Subject’s confidence rating plotted against their angle error.**

**S15 Fig. Subject’s confidence rating plotted against target location.**

**S16 Fig. Subject’s confidence rating plotted against the posterior variance.**

**S16 Fig. Subject’s confidence rating plotted against the posterior variance.**

**S1 Table. Parameters, their ranges and values, in the different models.**

## Acknowledgments

The Authors are grateful to Joshua Dean Garren for helping during data collection and Tapas Jaywant Arakeri for helpful discussions. Funding: This research was supported a grant from the National Science Foundation (NSF BCS-1630296 to PI Ekstrom).

## Supplementary Material

### 1 Computational Models

We built four models of the task which integrate the visual feedback in different ways. The simplest of these is the Path Integration model. This model ignores visual feedback completely and bases its estimate on pure path integration. As shown in the Results section, this model describes the behavior of at least one participant quite well and was used for all participants to fit data from the No Feedback condition.

The second model is the Kalman Filter model. This model integrates the visual feedback using the equations of the Kalman filter [55], which performs optimal cue combination under the assumption that the feedback error is Gaussian (which is not the case in our task because the feedback is sometimes sampled from a uniform distribution, see Equation 1).

The third and fourth models extend the Kalman filter to better match the actual generative process of the experiment. These models assume that the feedback can be misleading and take this possibility into account by computing the probability that the feedback is ‘true’ (i.e. comes from the Gaussian distribution, *p*(true|*f*)) and false (i.e. comes from the uniform distribution, *p*(false|*f*). The Cue Combination model, averages over this probability to form its estimate of heading. Conversely, the Hybrid model, samples from this probability, incorporating feedback just like the Kalman filter with probability *p*(true|*f*) and ignoring the feedback with probability *p*(false|*f*). In the following sections we develop each of these models in detail.

#### 1.1 Path Integration model

We begin by modeling the case in which feedback is either absent (as in the No Feedback condition) or ignored (as in some participants). In this case, the estimate of heading is based entirely on path integration of vestibular cues. To make a response, i.e. to decide when to stop turning, we assume that participants compare their heading angle estimate, computed by path integration, with their memory of the target angle. Thus, the Path Integration model can be thought of as comprising two processes: a path integration process and a target comparison process (Fig. S3).

##### Path integration

In the encoding phase of the task, participants are guided through an initial turn of −*α* degrees to face heading angle, *θ*_0_. In the retieval phase, they must then undo this rotation without visual feedback to return to *θ*_0_ + *α*. For simplicity, and without loss of generality, we take the initial head direction on each trial to be *θ*_0_ = 0°.

As they turn, we assume that participants receive vestibular cues about their angular velocity. For simplicity we model this process in discrete time, although the extension to continuous time is straightforward. We assume that on each time step *t* of the turn they receive a biased and noisy measure of their angular velocity, *d*_*t*_, which is related to their true angular velocity, *δ*_*t*_ by

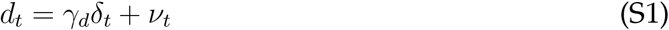

where *γ*_*d*_ denotes the gain on the velocity signal, which contributes to systematic under- or over-estimation of angular velocity. *ν*_*t*_ is zero-mean Gaussian noise with variance that increases in proportion to the magnitude of the angular velocity, |*δ*_*t*_|, representing a kind of Weber–Fechner law behavior [10],

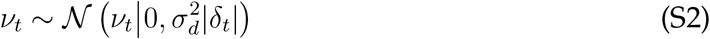

where 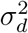 is a constant that determines the relationship between noise and angular velocity.

Next we assume that participants use this noisy velocity information to compute a probability distribution over their current heading angle.

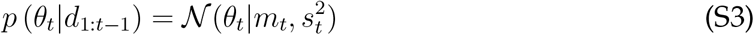

where the mean of the distribution is given by

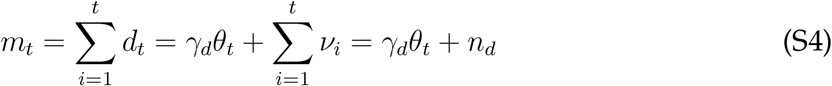

where 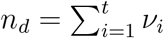. The variance of the distribution is given by

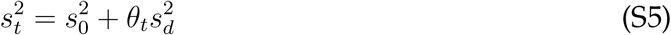

where 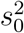 is the participant’s initial uncertainty in their location and 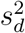 is the participant’s estimate of the variance of noise in their own vestibular system.

Strictly speaking, the behavior of the Path Integration model only depends on the mean, *m*_*t*_, and we do not need to make the assumption that participants compute a full distribution over possible heading. However, as we shall see, computing both the mean and variance of *p* (*θ*_*t*_|*d*_1:*t*−1_) will be necessary for the models that incorporate feedback.

**Fig S1.**
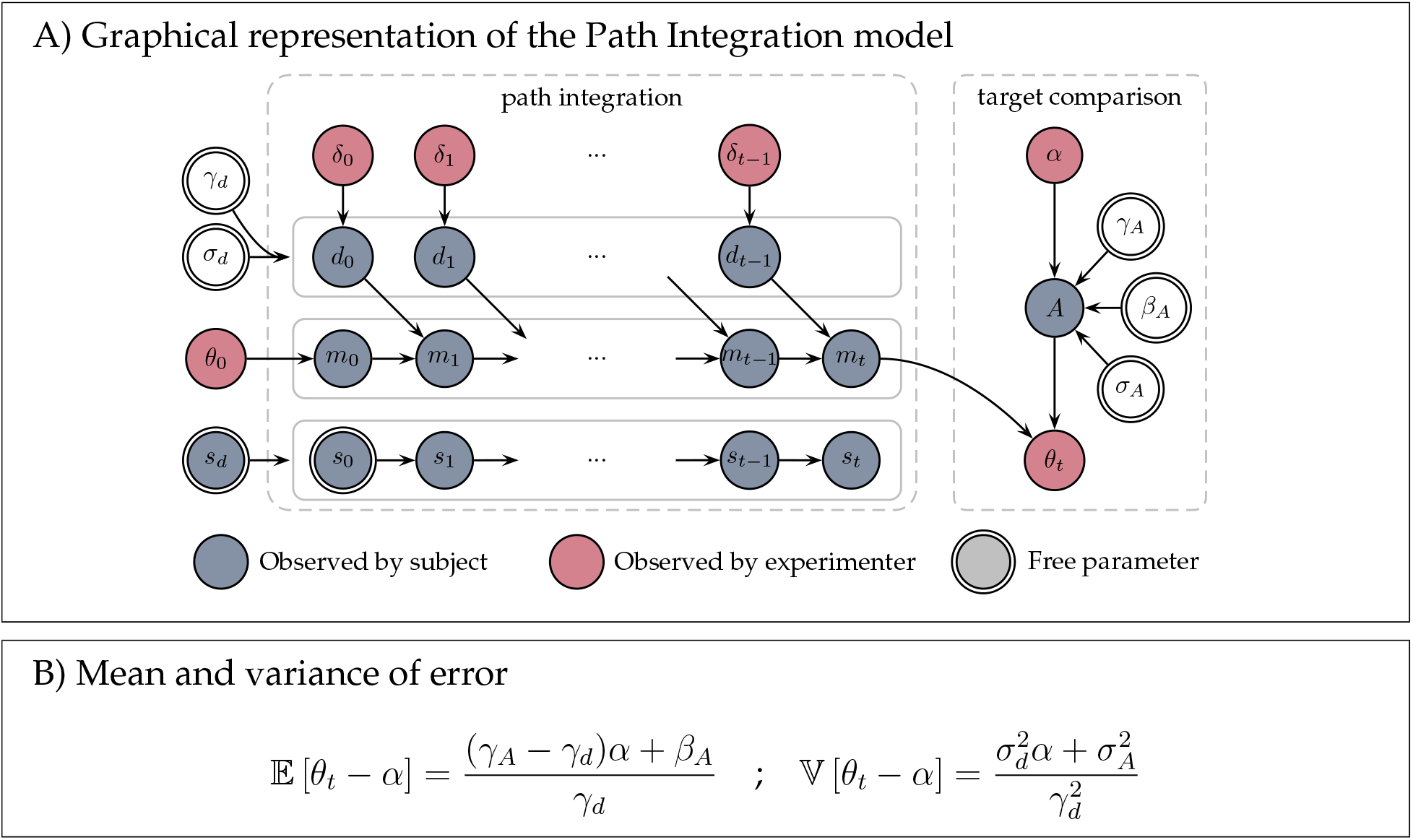
The Path Integration model. (A) The model comprises two processes: path integration and target comparison. In the path integration process the models’ estimate of heading angle, *m*_*t*_ (and corresponding uncertainty in this mean, *s*_*t*_) is computed by integrating biased and noisy velocity information, *d*_*t*_, over time. In the target comparison process, this estimate of heading is compared with the models’ biased and noisy memory of the target angle, *A*, to decide when to stop at measured angle *θ*_*t*_. Blue nodes correspond to variables that are ‘observed’ by the model (i.e. the participant) and can be used to compute the response. Red nodes correspond to variables that are observed by the experimenter and are the measurements we use to analyze behavior. White nodes correspond to parameters that are unobserved (by either the participant or the experimenter) describing imperfections in the coding of velocity (*γ*_*d*_, *σ*_*d*_) and target (*γ*_*A*_, *β*_*A*_, *σ*_*A*_). Free parameters are denoted by a double line. To further distinguish between variables available to the participant and those that are not, we write variables available to the participant with Roman letters and variables that are not available to the participant with Greek letters. (B) The model predicts that both the mean and variance of the response error will be linear in target angle, *α*.

##### Target comparison

Estimating the current heading angle is not enough to complete the task. In addition participants have to remember the target angle and compare it to their current estimate of heading. As with the encoding of velocity, we assume that this memory encoding is a noisy and biased process such that the participant’s memory of the target angle is

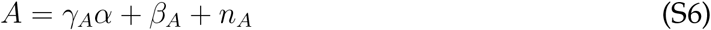

where *γ*_*A*_ and *β*_*A*_ are the gain and bias on the memory that leads to systematic over- or under-estimation of the target angle, and *n*_*A*_ is zero mean Gaussian noise with variance 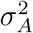. In the Supplementary Section 2 we show how this form of a gain and bias on the target angle can result from Bayesian decoding of a noisy memory with no gain or bias.

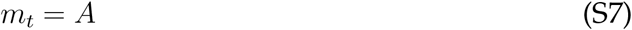

Substituting in the expressions for *m*_*t*_ and *A*, we get that the measured head angle when they stop, *θ*_*t*_, will satisfy

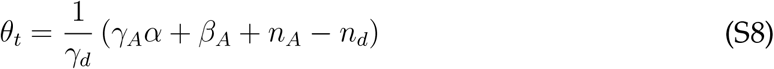

Because the noise terms (*n*_*A*_ and *n*_*d*_) are Gaussian, the measured error (*θ*_*t*_ − *α*) will also be Gaussian. Thus we can characterize the probability distribution over the measured error by its mean and variance

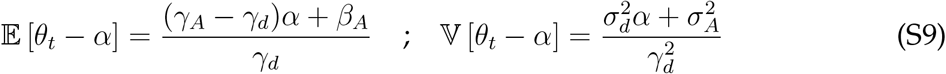

The Path Integration model technically has seven free parameters (*γ*_*d*_, *σ*_*d*_, *γ*_*A*_, *β*_*A*_, *σ*_*A*_, *s*_*d*_, *s*_0_). However, because the variance of the model’s estimate does not affect *θ*_*t*_, two of these parameters (*s*_*d*_ and *s*_0_) cannot be estimated. In addition, the remaining five parameters all appear as ratios with *γ*_*d*_ giving us four free parameters in the Path Integration model. In practice when fitting the Path Integration model we set *γ*_*d*_ = 1 and interpret the remaining parameters as ratios (e.g. *γ*_*A*_/*γ*_*d*_, Table S1).

#### 1.2 Kalman Filter model

Unlike the Path Integration model, which always ignores feedback, the Kalman Filter model always incorporates the visual feedback into its estimate of location. To model this process, we split the retrieval phase into four components: initial path integration, before the visual feedback is presented; feedback incorporation, when the feedback is presented; additional path integration, after the feedback is presented; and target comparison, to determine when to stop (Fig. S2).

##### Initial path integration

Initial path integration is identical to the Path Integration model. The model starts at a heading *θ*_0_ = 0 and integrates noisy angular velocity information over time to form an estimate of the mean, *m*_*t*_ and uncertainty, *s*_*t*_, over the current heading angle *θ*_*t*_.

##### Feedback incorporation

At time *t*_*f*_ the feedback, *f*, is presented. The Kalman Filter model assumes that the feedback can be noisy, but that it is always carries some information about the true heading angle 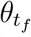. In particular, this model assumes that the feedback angle is sampled from a Gaussian distribution centered on the true heading angle, 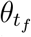, such that

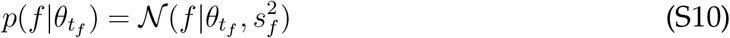

where 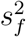 is the participant’s estimate of the variance of the feedback.

With this Gaussian assumption for the likelihood of the feedback, the Kalman Filter model then combines the feedback with the estimate from path integration via Bayes rule

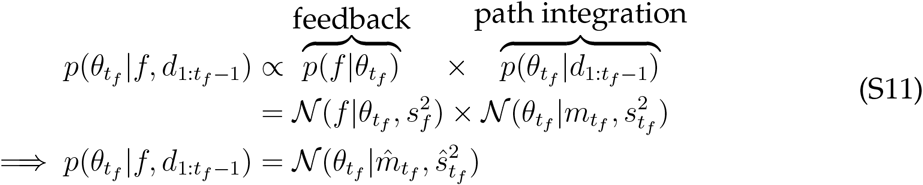

where the mean and variance of this posterior distribution over 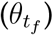 are given by

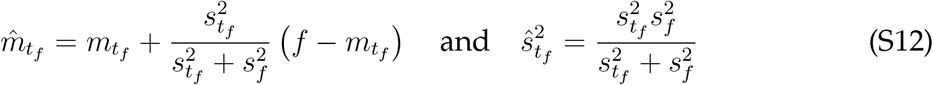

**Table S1.**
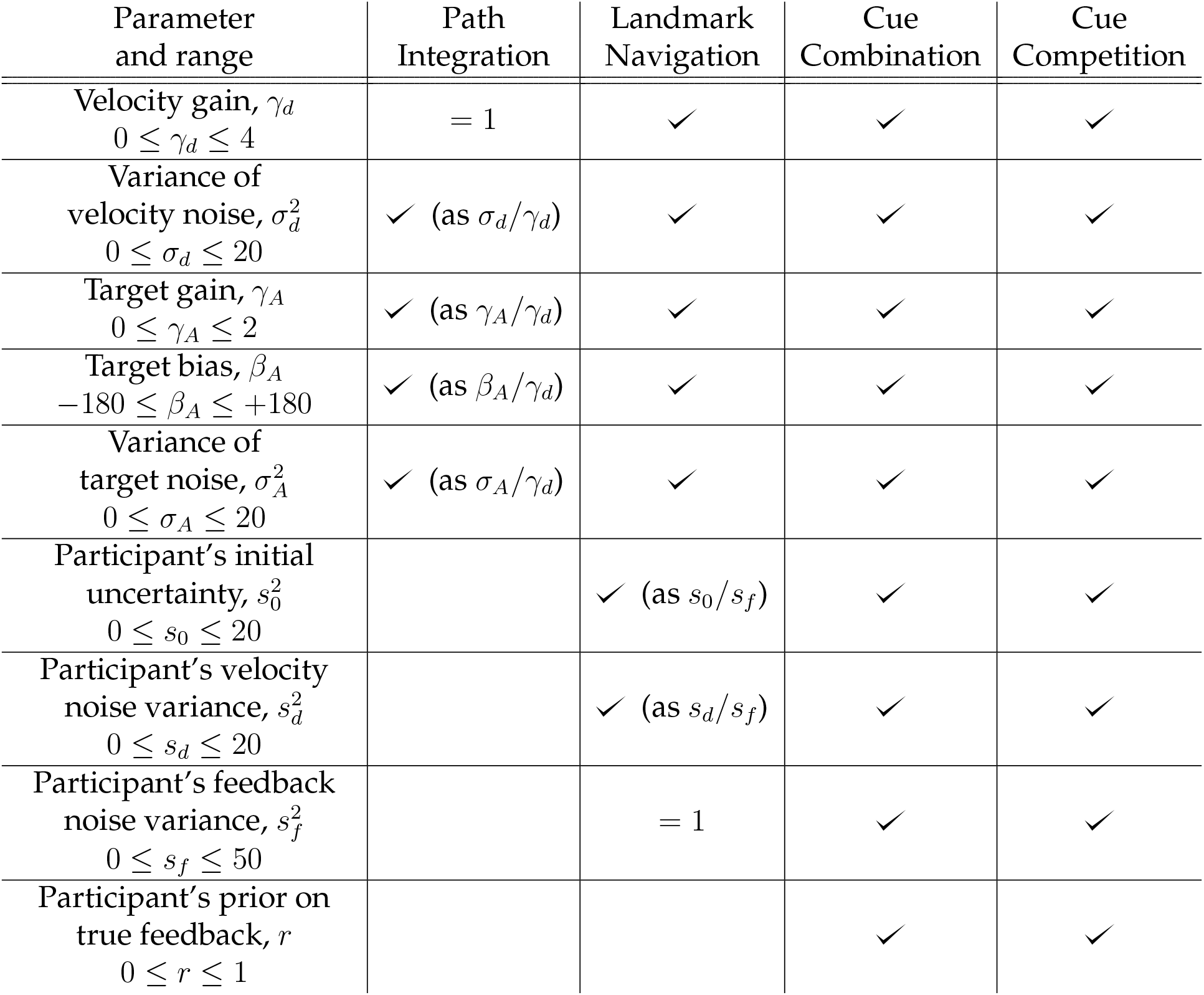
Parameters, their ranges and values, in the different models.

| Parameter and range | Path Integration | Landmark Navigation | Cue Combination | Cue Competition |
| --- | --- | --- | --- | --- |
| Velocity gain, $\gamma_d$<br>$0 \leq \gamma_d \leq 4$ | $= 1$ | ✓ | ✓ | ✓ |
| Variance of velocity noise, $\sigma_d^2$<br>$0 \leq \sigma_d \leq 20$ | ✓ (as $\sigma_d/\gamma_d$) | ✓ | ✓ | ✓ |
| Target gain, $\gamma_A$<br>$0 \leq \gamma_A \leq 2$ | ✓ (as $\gamma_A/\gamma_d$) | ✓ | ✓ | ✓ |
| Target bias, $\beta_A$<br>$-180 \leq \beta_A \leq +180$ | ✓ (as $\beta_A/\gamma_d$) | ✓ | ✓ | ✓ |
| Variance of target noise, $\sigma_A^2$<br>$0 \leq \sigma_A \leq 20$ | ✓ (as $\sigma_A/\gamma_d$) | ✓ | ✓ | ✓ |
| Participant's initial uncertainty, $s_0^2$<br>$0 \leq s_0 \leq 20$ | | ✓ (as $s_0/s_f$) | ✓ | ✓ |
| Participant's velocity noise variance, $s_d^2$<br>$0 \leq s_d \leq 20$ | | ✓ (as $s_d/s_f$) | ✓ | ✓ |
| Participant's feedback noise variance, $s_f^2$<br>$0 \leq s_f \leq 50$ | | $= 1$ | ✓ | ✓ |
| Participant's prior on true feedback, $r$<br>$0 \leq r \leq 1$ | | | ✓ | ✓ |

**Fig S2.**
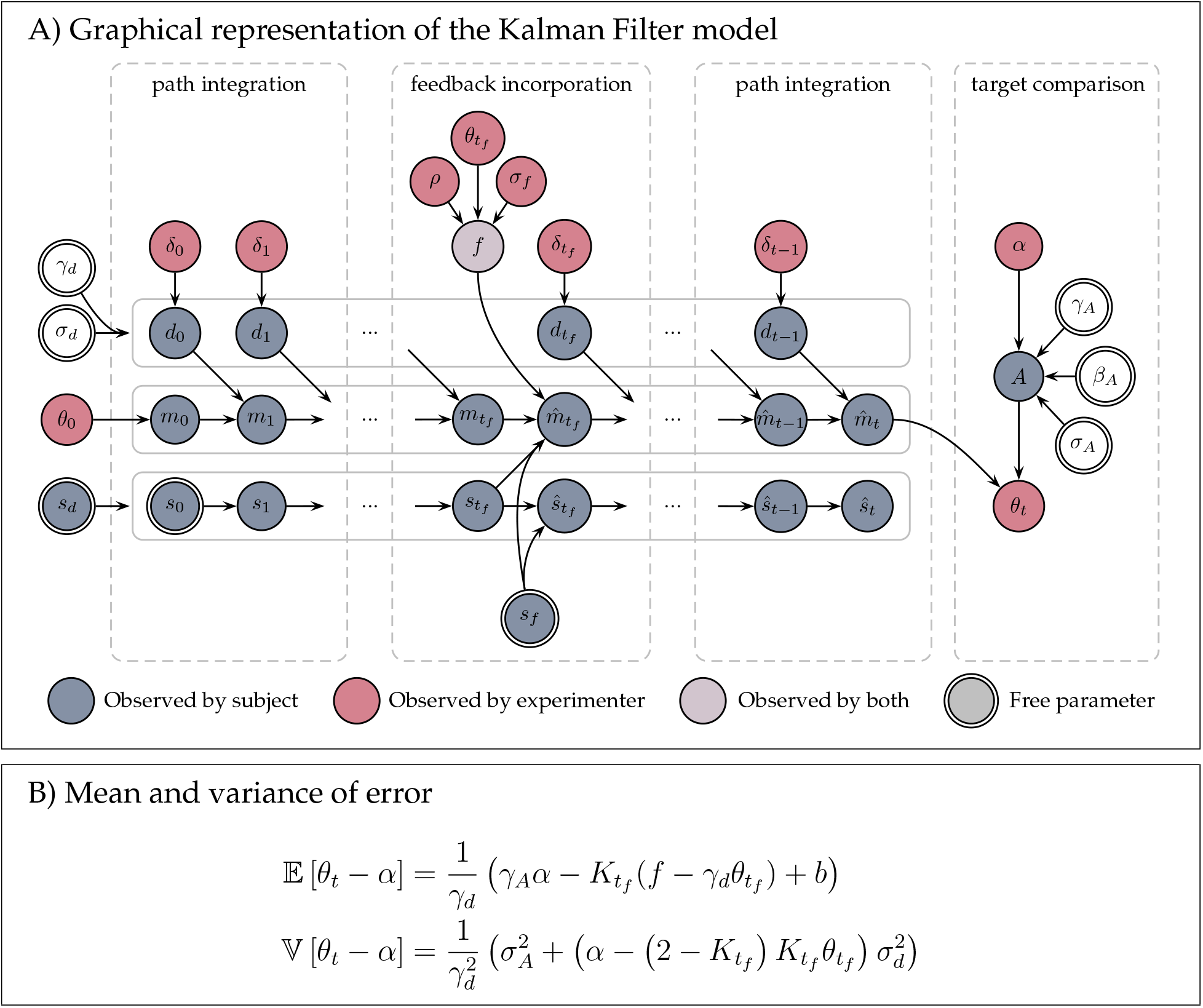
Graphical representation of the Kalman Filter model. (A) The model comprises four processes: initial path integration, feedback incorporation, final path integration, and target comparison. The initial path integration process proceeds exactly as in the Path Integration model, estimating heading angle *m*_*t*_ from noisy velocity information *d*. In the feedback incorporation process, the path integration estimate is combined with the feedback *f* to form a combined estimate 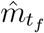. In the final path integration process, the model incorporates the new noisy velocity information to update the combined estimate of heading. Finally, in the target comparison process, the combined estimate is compared with the remembered target angle *A* to generate the response *θ*_*t*_. (B) Expressions for the mean and variance of the measured response error. Of note is that the mean is linear in both the target angle and the prediction error 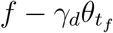.

Note that the feedback updates the mean according to the prediction error 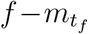 weighted by the ‘Kalman gain’

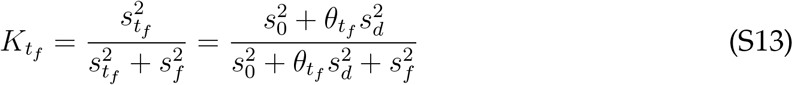

The Kalman gain captures the influence of the prediction error on the estimate of heading angle. The more certain the model is that the feedback is accurate (i.e. smaller *s*_*f*_) the closer the Kalman gain is to 1 and the larger the effect of the feedback. Conversely, the more certain the model is in its path integration estimate (i.e. smaller 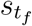) the closer the Kalman gain is to 0 and the smaller the effect of the feedback.

##### Additional path integration

After the feedback has been incorporated, the model continues path integration using noisy velocity information. Thus the estimate of the mean continues to update as:

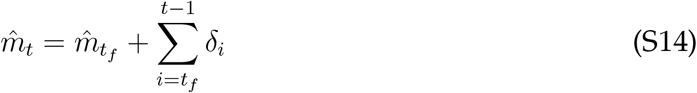

Substituting in the expressions for 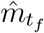, *d*_*i*_, and 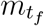 we get

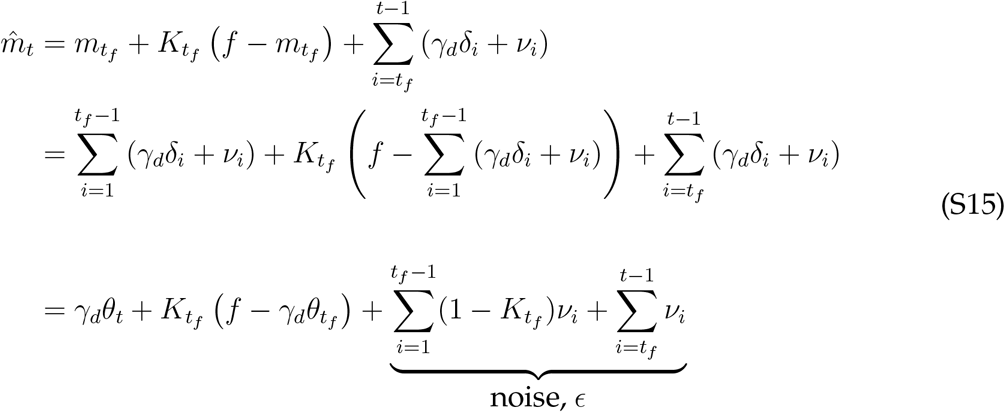

##### Target comparison

Finally, the response is determined as the point at which 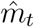 is equal to the noisy target angle, *A*

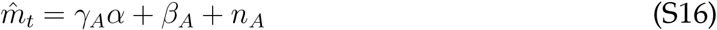

Substituting in the expression for 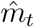 and rearranging for the response angle gives

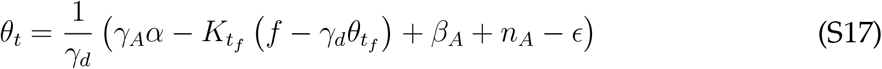

This implies that the distribution of errors (*θ*_*t*_ − *α*) is Gaussian with a mean given by

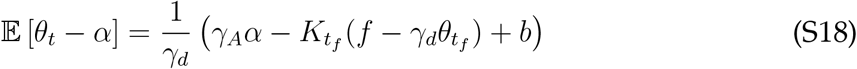

and a variance given by

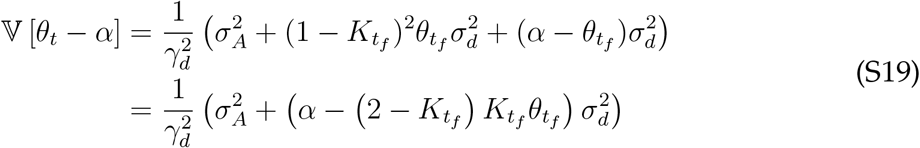

Note that, because 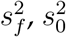, and 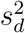 appear as part of a ratio in the equation for 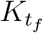, only two out of the three of these can be estimated from data. Thus, of the eight free parameters in the Kalman Filter model (*γ*_*d*_, *σ*_*d*_, *γ*_*A*_, *β*_*A*_, *σ*_*A*_, *s*_0_, *s*_*d*_, and *s*_*f*_), only seven can be estimated from the data. In practice, when fitting this model we set 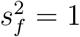 and interpret the participant’s initial uncertainty and estimate of velocity noise as ratios, *s*_0_/*s*_*f*_ and *s*_*d*_/*s*_*f*_ respectively (Table S1).

#### 1.3 Cue Combination model

The Cue Combination model takes into account the possibility that the feedback will be misleading. To do this it computes a mixture distribution over heading angle with one component of the mixture assuming that the feedback is false and the other that the feedback is true. These two components are weighted according to the computed probability that the feedback is either false or true.

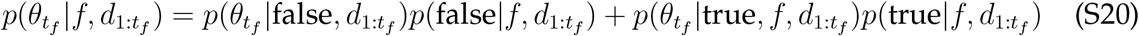

where 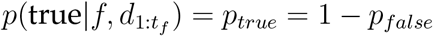 is the probability that the feedback is true given the noisy velocity cues seen so far. Thus the Cue Combination model requires two steps to incorporate the feedback: first compute *p*_*true*_ and second average over *p*_*true*_ to determine how much to take the feedback into account.

##### Computing *p*_*true*_

Using Bayes rule, we can write the probability that the feedback is true, *p*_*true*_ as

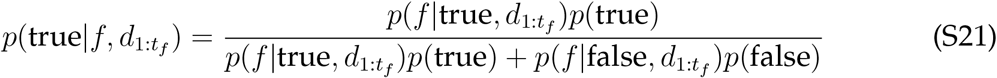

where 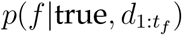 is the likelihood of the feedback assuming it is true, 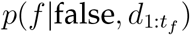 is the likelihood of the feedback assuming it is false, and *p*(true) = 1 − *p*(false) is the participant’s estimate of the prior probability that the feedback is true.

**Fig S3.**
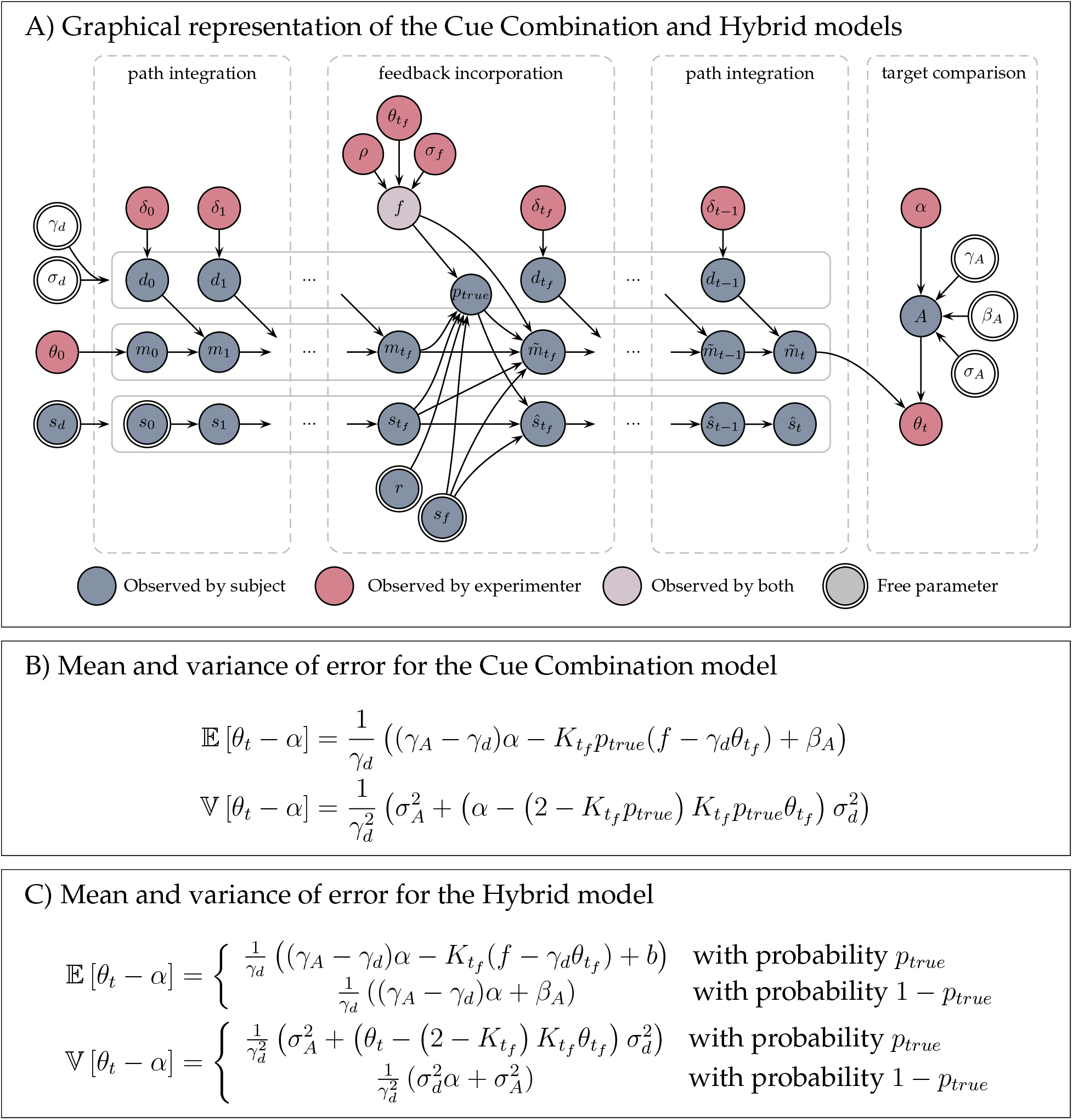
The Cue Combination and Hybrid models. (A) Graphical representations of the parameters in the Cue Combination and Hybrid models. In the feedback incorporation stage, both models compute the probability that the feedback is true, *p*_*true*_. They then use this probability to modulate the effect of feedback on their estimate of heading — the Cue Combination model by averaging over *p*_*true*_, the Hybrid model with sampling from *p*_*true*_. (B) The mean and variance of the measured error for the Cue Combination model are linear in the target angle, *α*, but non-linear in the prediction error, 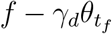 because *p*_*true*_ is non-linear in the prediction error. (C) The response distribution for the Hybrid model is a mixture of two Gaussians.

The likelihood of the feedback given that it is true, 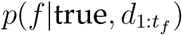, can be computed by marginalizing over the estimate of heading direction as

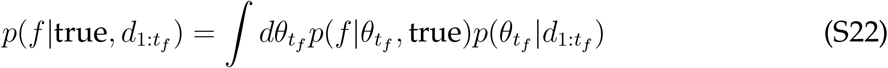

where 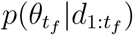 is the heading angle distribution computed by path integration and 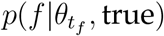 is the likelihood of the feedback given the heading angle, which we assume to be Gaussian; i.e.,

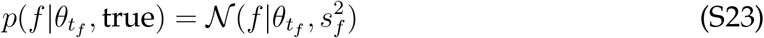

This implies that 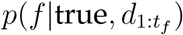 is the convolution of two Gaussians, which is itself another Gaussian

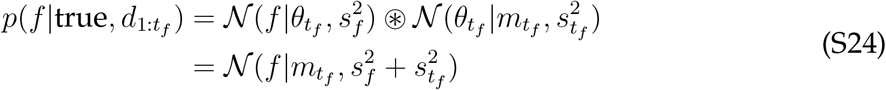

The likelihood of the feedback given that it is false, 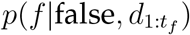, is simply a uniform distribution over *f*

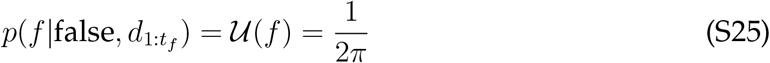

Finally, we define the participant’s estimate of the prior probability that the feedback is true as a free parameter *p*(true) = 1 − *p*(false) = *r*.

Putting it all together gives the following expression for *p*_*true*_

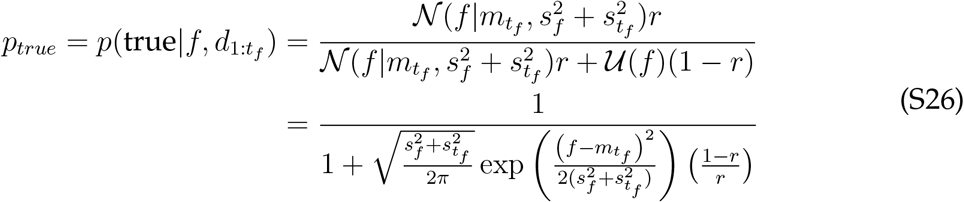

Note, that *p*_*true*_ depends on the square of the prediction error 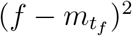 such that when the prediction error has a large magnitude, *p*_*true*_ is small. Unfortunately this dependence requires an approximation before we can use it for model fitting. The reason is that 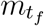 is not observed by the experimenter, only 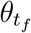. In addition, because the dependence of *p*_*true*_ on 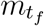 is non-linear it is not easy to average over this exactly. Instead we approximate 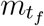 in the expression for *p*_*true*_ with its average, 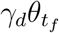, thus our approximate expression for *p*_*true*_ becomes

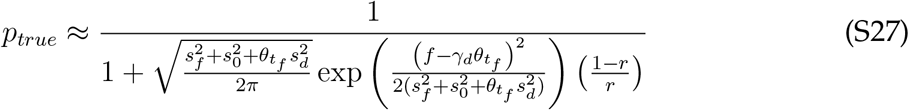

Note that *s*_0_, *s*_*d*_, and *s*_*f*_ do *not* appear as a ratio in the expression for *p*_*true*_. This implies that, unlike the Kalman Filter model, all three of these parameters can be estimated from the data.

##### Cue Combination over *p*_*true*_

The Cue Combination model incorporates feedback by computing the mixture distribution

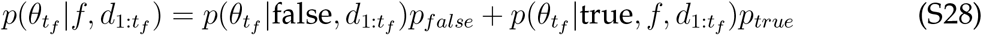

This sum is the weighted sum of the distributions from the Path Integration model, 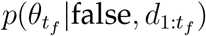, and the Kalman Filter model, 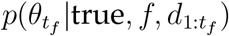. Thus, at time *t*, the mean of the combined heading angle distribution is simply the weighted sum of the Path Integration estimates and the Kalman Filter estimates; i.e.,

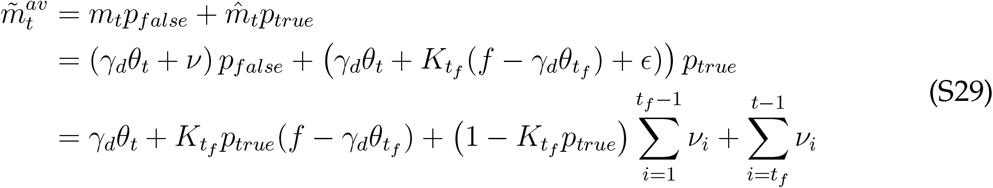

Computing the variance of 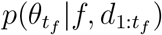 is a little more involved. However, because this variance is unrelated to the measured behavior, we ignore it.

##### Target comparison

As with the other models, we assume that participants stop turning when their estimate of the mean heading angle matches their noisy memory of the target, i.e. when

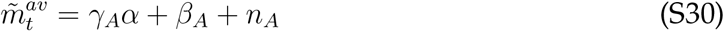

Rearranging for the measured response angle, *θ*_*t*_, gives

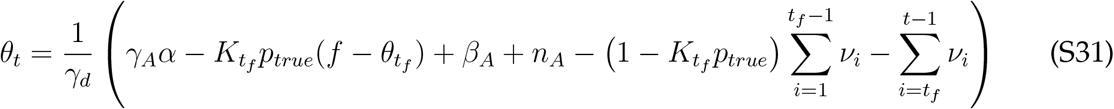

Which implies that the error follows a Gaussian distribution with mean and variance given by

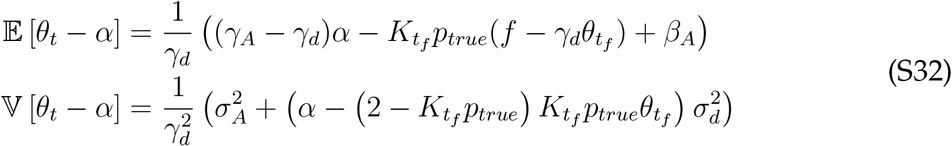

The Cue Combination model has nine free parameters (Table S1), all of which can be estimated from the data.

#### 1.4 Hybrid model

Instead of averaging over the possibility that the feedback is true or false, the Hybrid model makes a decision to either incorporate the feedback (in the same way as the Kalman Filter model) or ignore it (in the same way as the Path Integration model) (Fig. S3). We assume that the model makes this decision according to *p*_*true*_, by sampling from the distribution over the veracity of the feedback. Thus with probability *p*_*true*_, this model behaves exactly like the Kalman Filter model, with

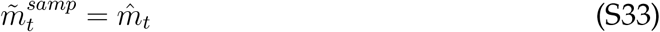

and with probability *p*_*false*_ = 1 − *p*_*true*_ this model behaves exactly like the Path Integration model with

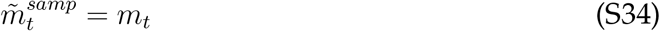

This implies that the distribution of errors is a mixture of two Gaussians, such that with probability *p*_*true*_, the mean and variance of the response error are

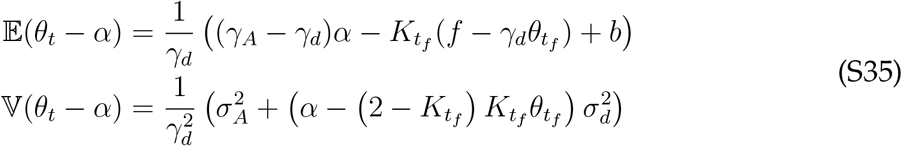

and with probability *p*_*false*_ = 1 − *p*_*true*_, the mean and variance of the response error are

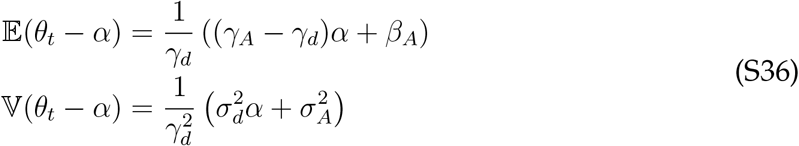

Like the Cue Combination model, the Hybrid model has nine free parameters (Table S1) all of which can be estimated from the data.

### 2 Bayesian decoding of target position

In the main text we assumed the following form for the biased and noisy memory of the target

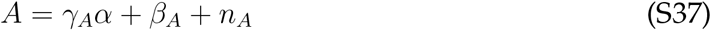

where *γ*_*A*_ and *β*_*A*_ are the gain and bias on the memory, which lead to systematic over- and under-estimation of the target angle, and *n*_*A*_ is zero mean Gaussian noise with variance 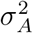.

Here we show how this expression can be related to Bayesian decoding of a noisy, but otherwise unbiased target angle

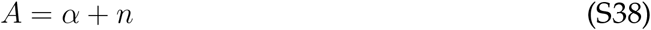

In particular, we assume that participants are aware that their memory is imperfect and can, to some degree correct for this noise by incorporating prior knowledge about possible *α* angles. That is, participants use Bayesian inference compute a posterior over *α* given *A* as

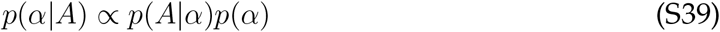

Assuming both the prior and likelihood are Gaussian such that

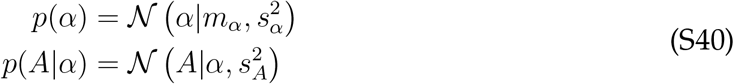

where *m*_*α*_ is the participant’s estimate of the mean of the prior distribution, 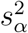 is their estimate of the variance of the prior, and 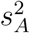 is the their approximation to the variance of the memory noise (i.e. their estimate of the variance of *n*_*A*_).

Substituting these expressions for the likelihood and prior into Equation S39 implies that the posterior over target angle is also a Gaussian with a mean and variance given by Note that the mean of this distribution is

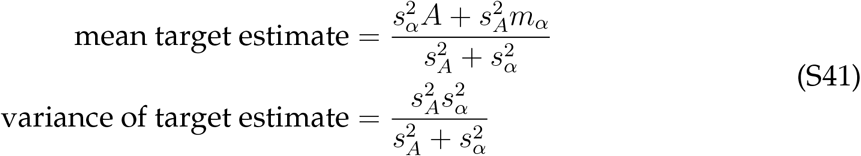

Note that the expression for the mean can be further related to the target by substituting *A* = *α* + *n* giving

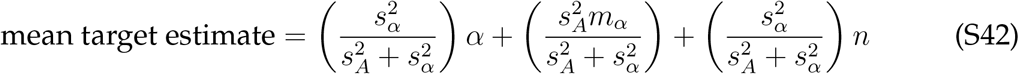

Comparing this expression for the mean with Equation S37 we can make the identifications

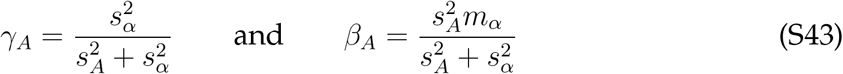

### 3 Fitting simulated data

We tested the validity of our model fitting procedure by fitting simulated data. This allowed us to determine whether data generated by a given model would be best fit by that model (model recovery) and whether the parameters used to generate the data could be recovered by the fitting process (parameter recovery). Matlab code to reproduce these analyses can be found on GitHub (https://github.com/sharootonian/HeadingDirection).

#### 3.1 Simulated data

Simulated data for each model were generated by simulating the models using the generative processes described in the main text. To ensure that the parameter values used to simulate data were in a reasonable range, we used the parameter values fit to the participants’ behavior to generate parameters for the simulations. In particular, for each simulated participant we sampled each parameter randomly from the values fit to the participants. Thus, the first simulated participant could have *γ*_*d*_ from participant number 5, *σ*_*d*_ from participant number 27, and so on. In this way we ensured that the simulation parameters were in a reasonable range, but removed any correlations between the parameters in the simulation. This latter point was important for testing whether the fitting procedure induced correlations between the parameters. In all we simulated behavior from 30 participants per model (total 120 simulated participants) on the same (but scrambled) set of trials seen by real participants in the experiment. Thus the simulated data we obtained had the same number of trials as the real data set, but four times the number of participants (30 per model).

#### 3.2 Fitting simulated data

We fit the simulated data using the same procedure used to fit the real data in the main paper. This allowed us to compute the best fitting parameter values and a BIC score for each

#### 3.3 Model recovery

We tested the ability of the model to identify the generating model by fitting all 120 simulated data sets with all four models. Using the BIC scores for each participant, we then computed the ‘confusion matrix’ [56] as the fraction of times that data generated by model *X* was best fit by model *Y*, *p*(fit = *Y*|sim = *X*) (Fig. S4A). In a perfect world this matrix would be the identity matrix indicating that data generated by model *X* is always best fit by model *X*. In practice, limitations in the experiment design and fitting procedure often cause these matrices to be non-diagonal as is the case here. Nevertheless, for every model, more than 50% of the data sets generated by model *X* are best fit by model *X*.

To further help interpret the model recovery data, we also computed the ‘inversion matrix’ [56]. Unlike the confusion matrix, which approximates *p*(fit|sim), the inversion matrix approximates *p*(sim|fit). This we compute from the confusion matrix using Bayes rule

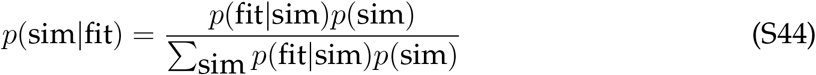

under the assumption that the prior on generating models *p*(sim) is uniform.

The inversion matrix more closely matches the inference process we face when interpreting the model fitting data in the paper. That is, we observe which model best fits each subject and must infer the model that generated it. Again we see that model recovery is good, but not perfect, such that (for example) 74% of the time that a model is best fit with the Cue Competition model it was actually generated by the Cue Competition model.

**Fig S4.**
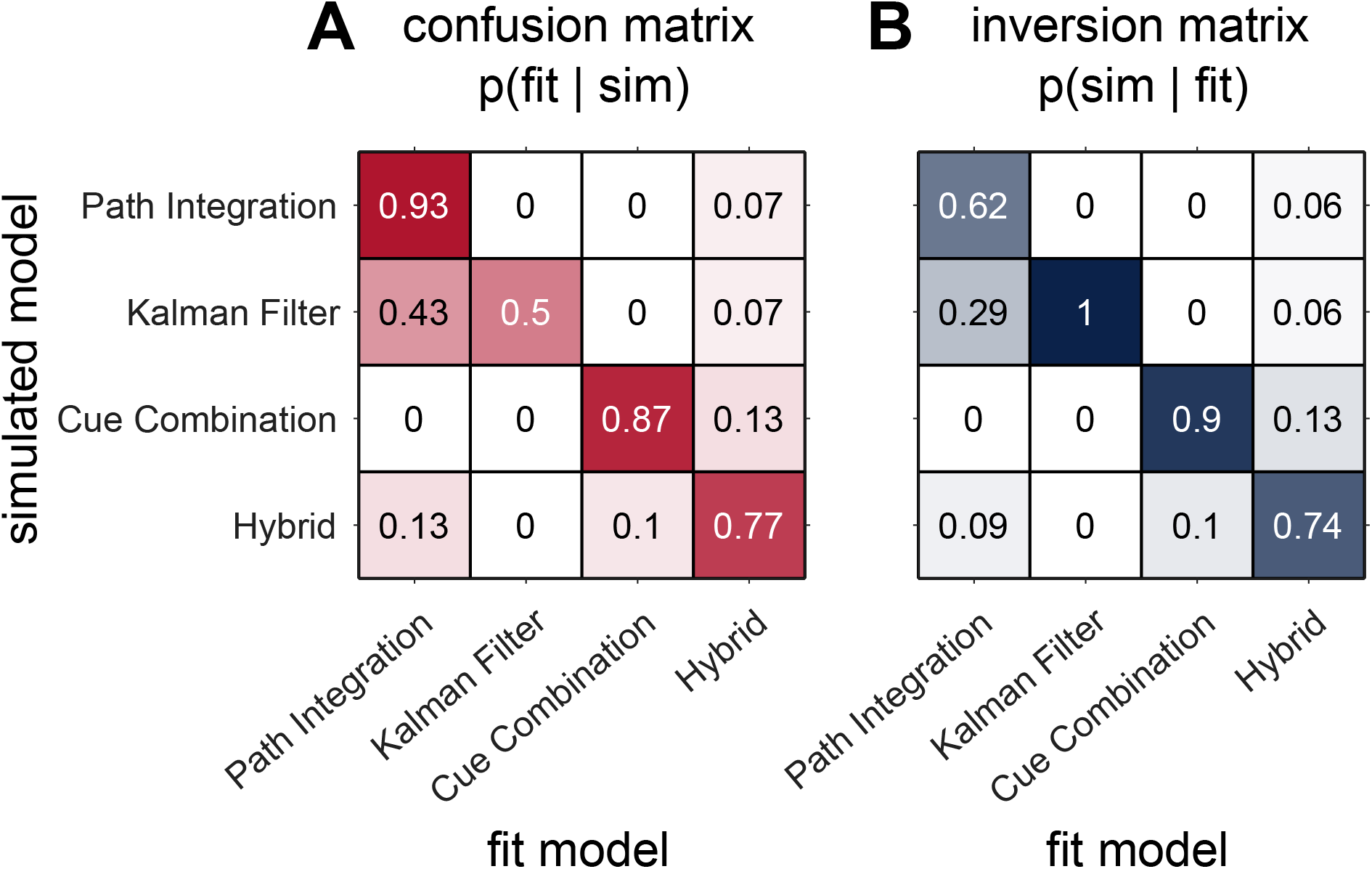
Confusion matrix showing the frequency with which data simulated according to each model is best fit by another model, *p*(fit model|simulated model)

#### 3.4 Parameter recovery

For the parameter recovery analysis we simulated and fit the data with the same model. First, we fit data from the No Feedback model on just the No Feedback trials S5. Parameter recovery is excellent int his case with no correlation between simulated and fit data falling below 0.84.

**Fig S5.**
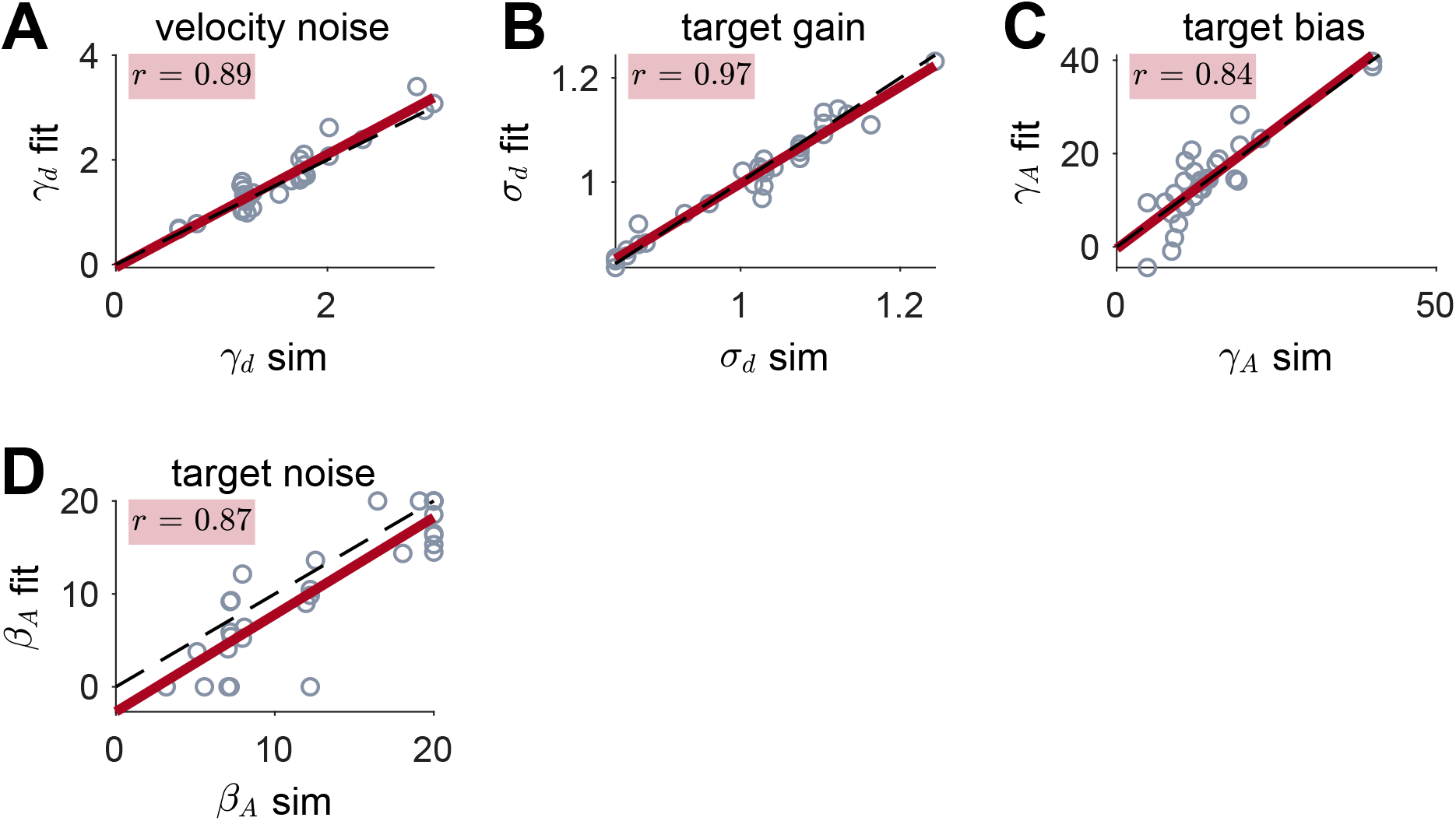
Parameter recovery for Path Integration model

Next we performed the same analysis for the Cue Competition model, this time fitting all trials, including both the No Feedback and Feedback conditions (Fig. S6). Again, parameter recovery is good for this model.

**Fig S6.**
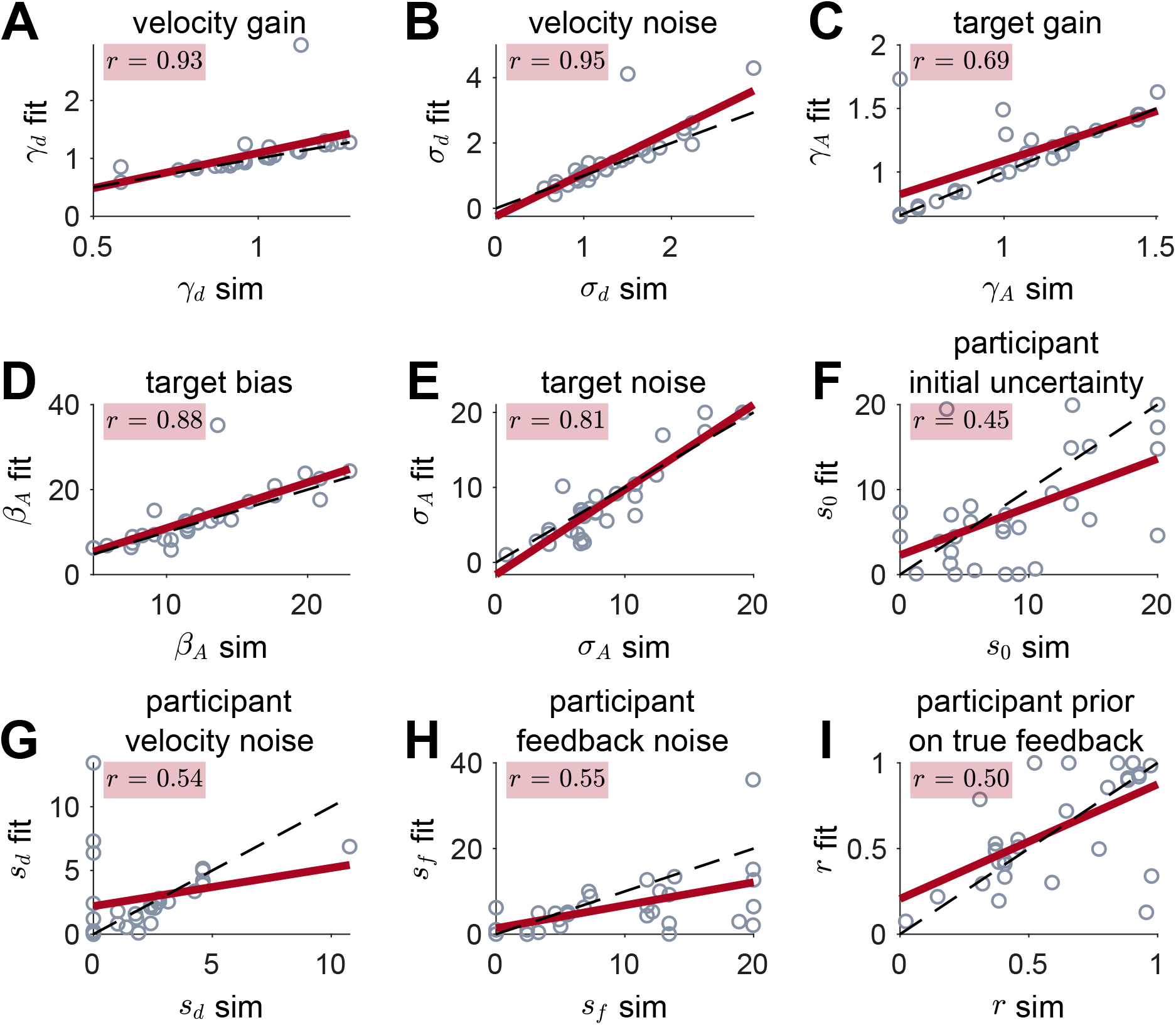
Parameter recovery for Hybrid model

Critically, the model fitting process did not introduce new correlations into the data set. In Fig. S7 we show this for the correlations between *γ*_*A*_ and *γ*_*d*_ and *σ*_*d*_.

**Fig S7.**
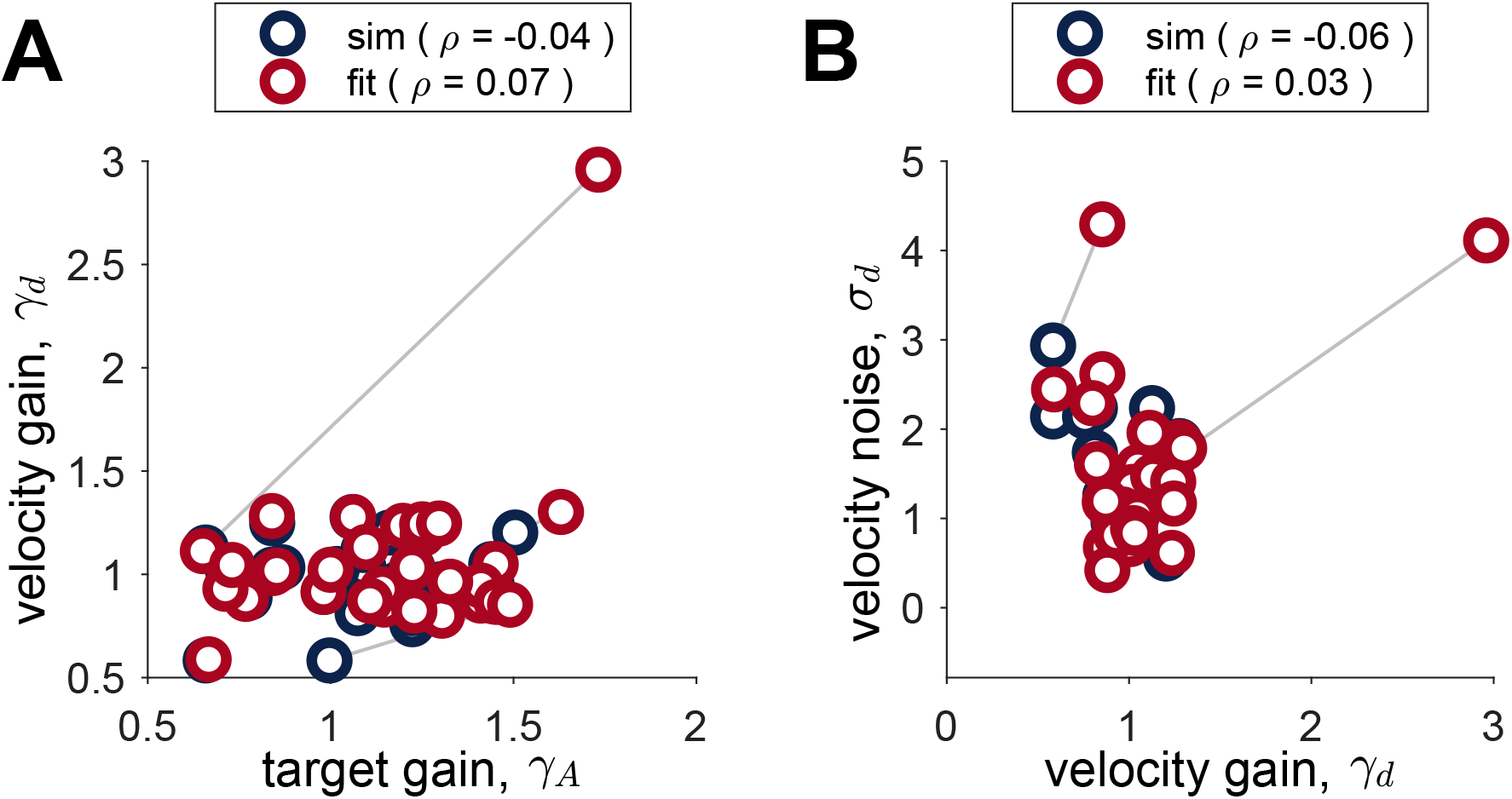
The fitting procedure does not induce correlations between parameters either for *γ*_*d*_ and *γ*_*A*_ (A) or *γ*_*d*_ and *σ*_*d*_ (B). When the simulation parameters (red) are uncorrelated, so are the fit parameters (blue). Gray lines connect simulated and fit parameters.

Finally, for completeness we performed parameter recovery for the remaining three models (No Feedback, Landmark Navigation, and Cue Combination model) using all trials (i.e. from the No Feedback as well as the Feedback condition). Parameter recovery was pretty good for all models Figures S8, S9, and S10.

**Fig S8.**
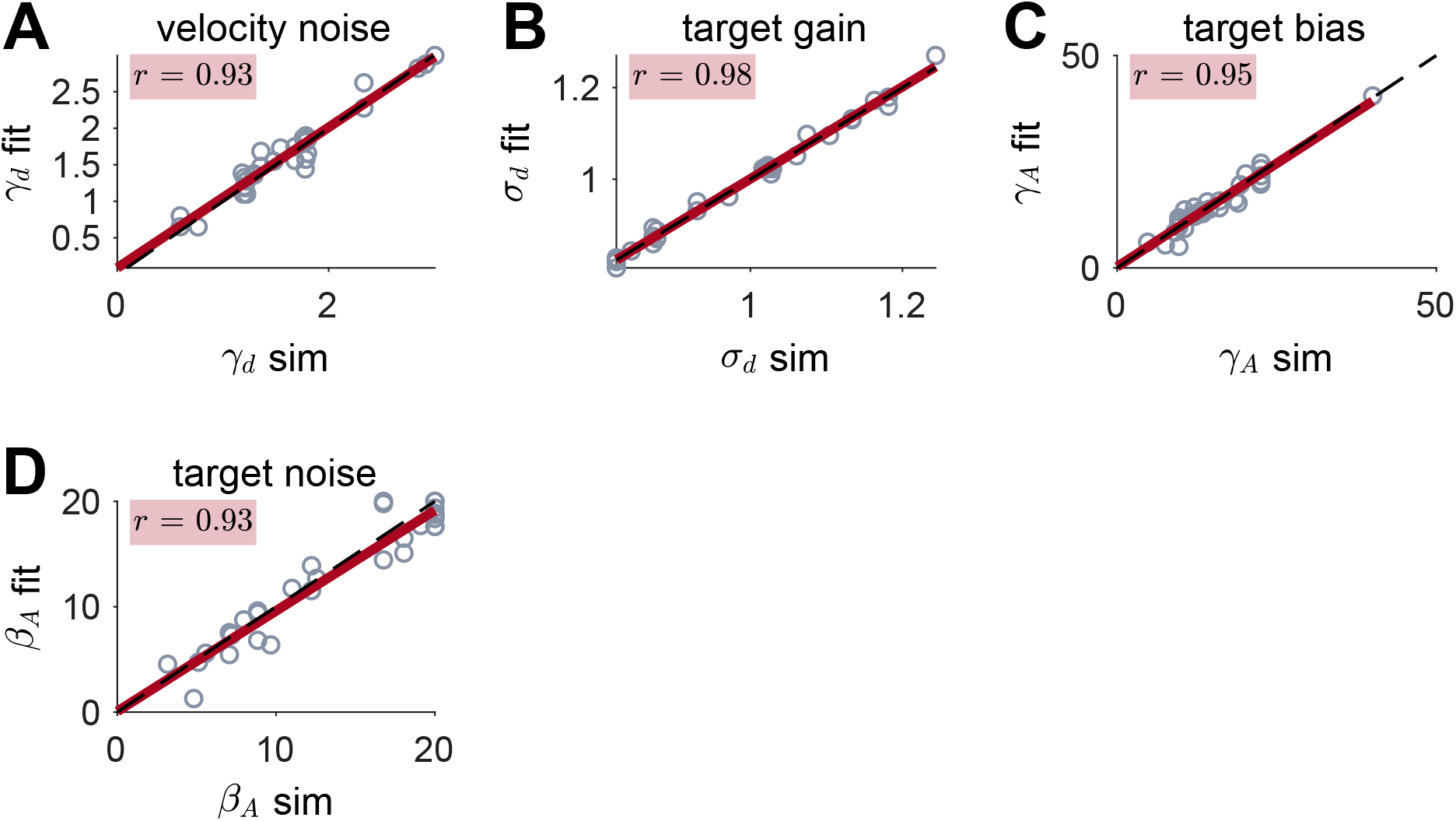
Parameter recovery for Path Integration model

**Fig S9.**
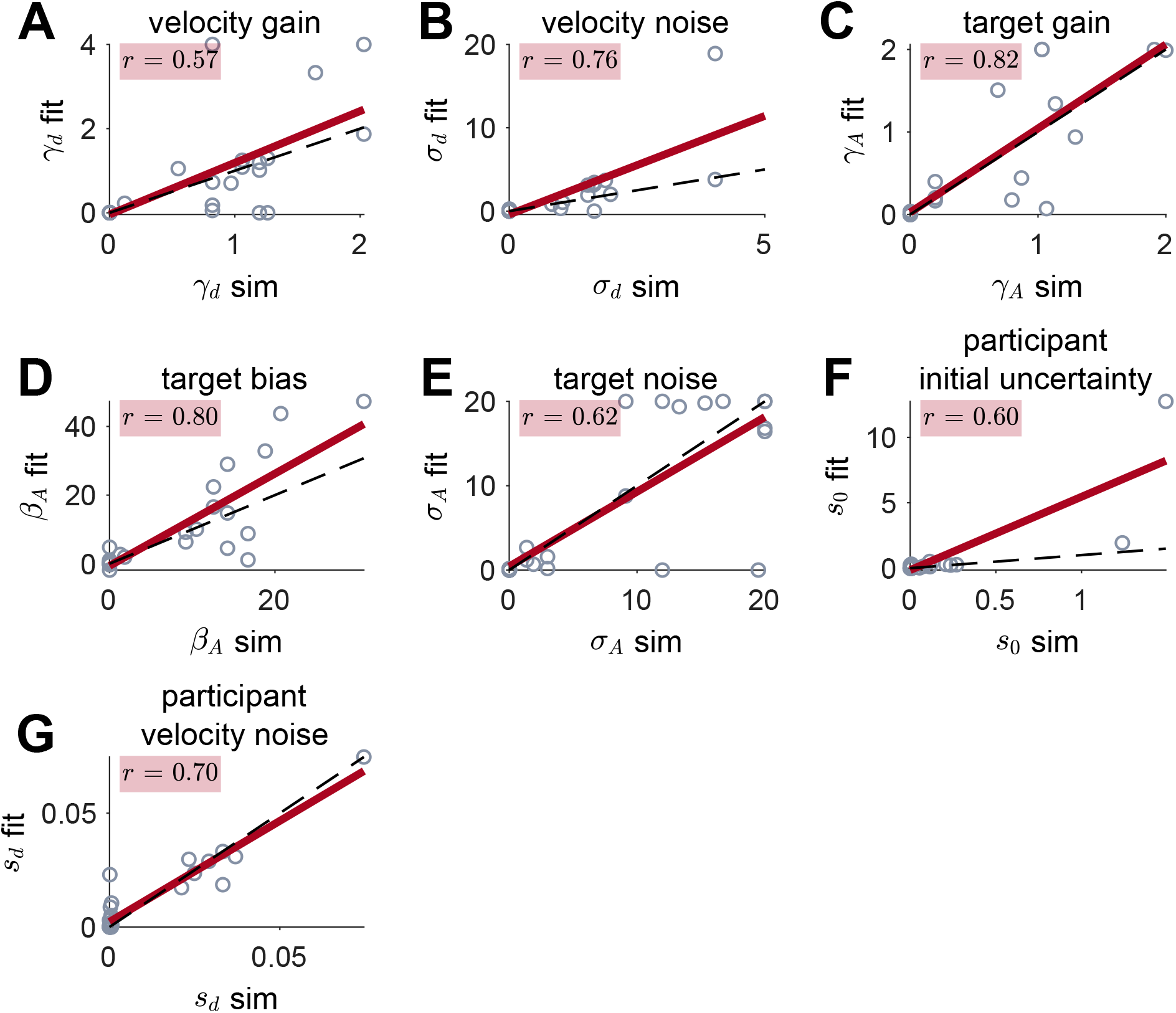
Parameter recovery for Kalman Filter model

### 4 Parameter values for the Hybrid model

Fit parameter values for the Hybrid model are shown in Fig. S11. Like the model-free measures of behavior, there was considerable variability in the parameter values across participants. Thus, while the group average of the velocity gain was close to 1 (mean *γ*_*d*_ = 1.01) individuals varied from systematically under-weighting velocity (*γ*_*d*_ < 1) to systematically over-weighting it (*γ*_*d*_ > 1). All participants exhibited noise in their velocity coding process, with *σ*_*d*_ = 1.38 on average. This latter result suggests that the variance of the uncertainty in location from path integration grows rapidly, at around 1.38 times the rotation angle. At this rate of growth, the noise in path integration will swamp the signal in less than one turn.

Similarly suboptimalities were observed in the coding of the target. Like the velocity gain, the group average of the target gain was close to 1 (mean *γ*_*A*_ = 1.04), there was considerable variation between people from systematic under-weighting (*γ*_*A*_ < 1) to systematic over-weighting of the target (*γ*_*A*_ > 1). In addition, as in the No Feedback condition, all participants were biased towards over-estimating the target (mean *β*_*A*_ = 13.0 degrees) and had considerable noise in the target coding process (mean *σ*_*A*_ = 7.5 degrees).

There were also considerable individual differences in participants’ inference parameters: *s*_0_, *s*_*d*_, *s*_*f*_, and *r*. Most participants underestimated the feedback noise (mean *s*_*f*_ = 8.68 versus the true value in the experiment of *σ*_*f*_ = 30°). Conversely, the group average of the prior probability was more accurate (mean *r* = 0.69 which is remarkably close to the true value of *ρ* = 0.7, although again there was considerable variability across the group.

The individual differences in *s*_0_, *s*_*d*_, and *s*_*f*_ lead to considerable variability in the Kalman gain across participants and (in some participants) across trials (Fig. 12). Some participants show almost no variation across trials (left and right sides of Fig. 12), while others show large variability across trials (middle participants in Fig. 12). This pattern can be explained by recalling the equation for Kalman gain

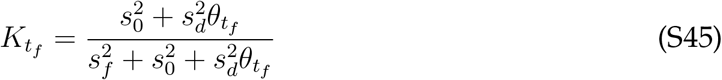

This equation implies that participants with both small and large noise in their path integration process (i.e. *s*_*d*_ ≈ 0 or *s*_*d*_ ≫ *s*_*f*_ and *s*_0_) will have approximately constant Kalman gain across trials. When the velocity noise is small (*s*_*d*_ ≈ 0),

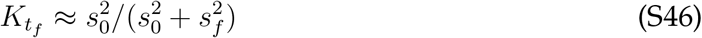

which is a constant between 0 and 1 depending on the ratio of participants initial uncertainty *s*_0_ to their estimate of feedback noise *s*_*f*_. This is the case for participants on the left hand side of Fig. 12.

When the velocity noise is large (*s*_*d*_ ≫ *s*_*f*_ and *s*_0_)

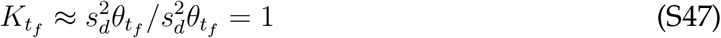

That is the Kalman gain is a constant with value equal to 1. This is the case for participants on the right hand side of Fig. 12.

**Fig S10.**
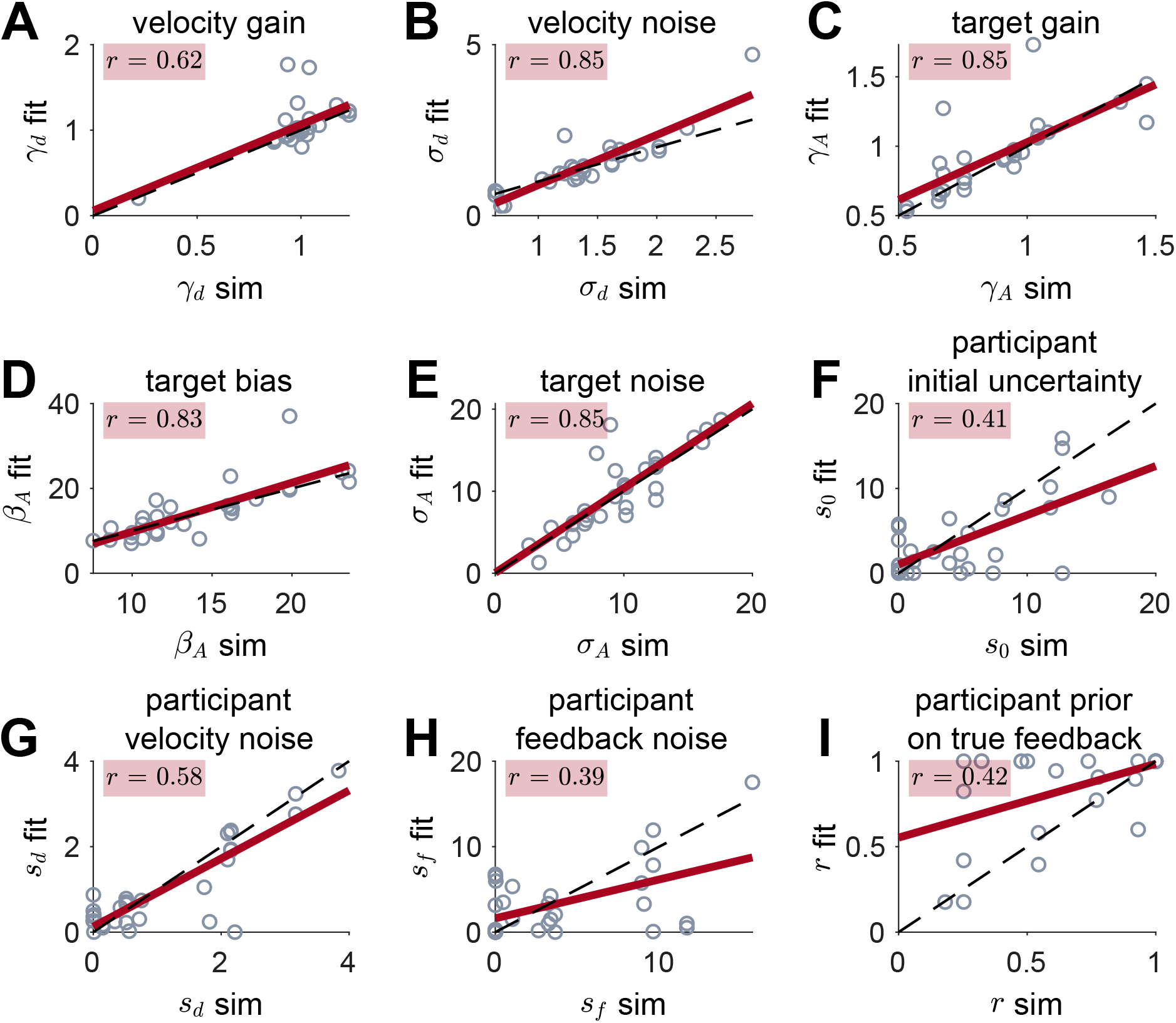
Parameter recovery for Cue Combination model

**Fig S11.**
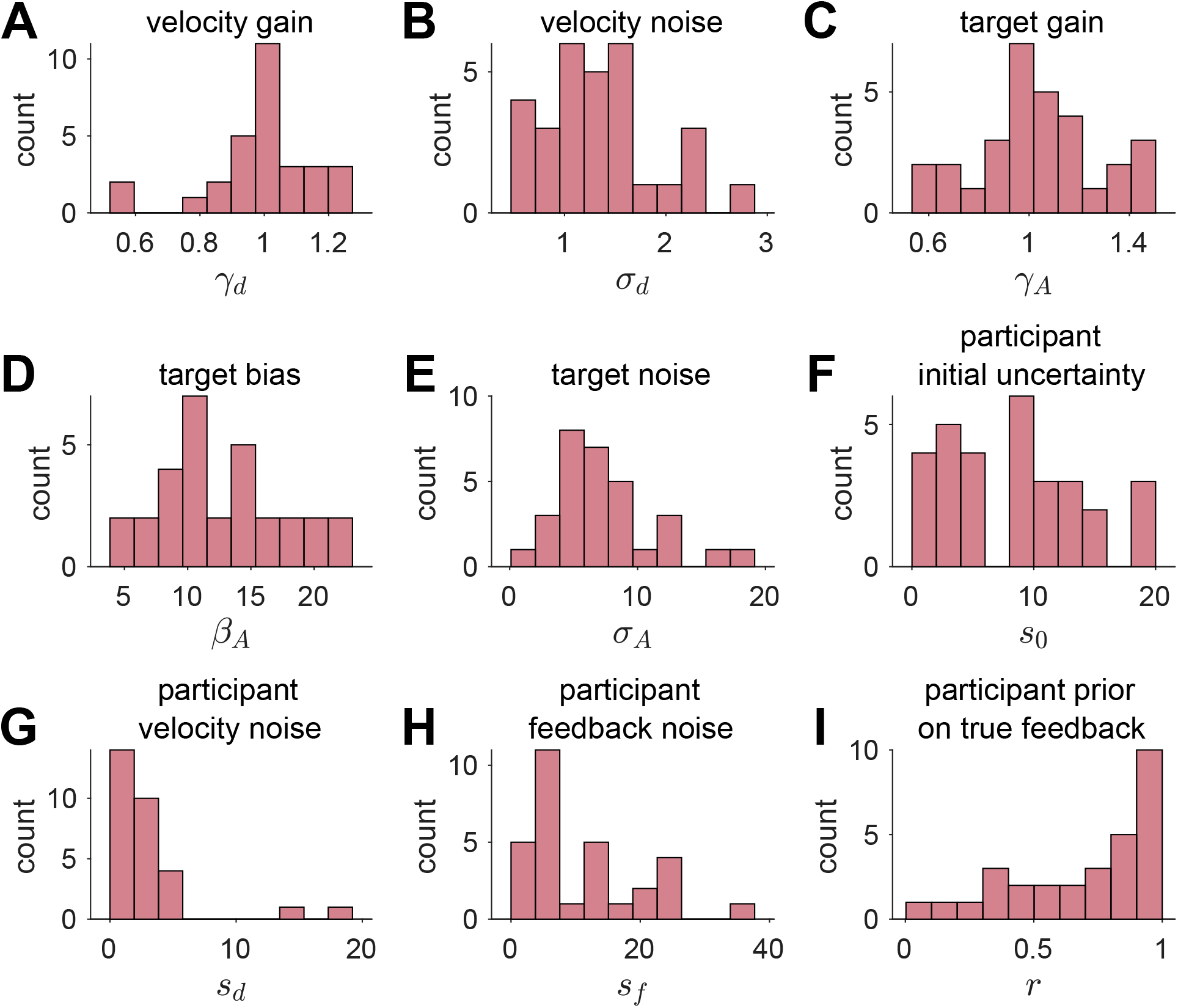
Best fit parameters for the Hybrid model

For participants whose velocity noise is intermediate in value, the uncertainty in their estimate of heading at the time of feedback is close to their estimate of the feedback noise giving them a Kalman gain between 0 and 1 that varies considerably depending on the exact value of the true heading angle at feedback 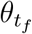.

#### 4.1 Correlations between parameters

Finally we consider the correlations between fit parameter values (Fig. S12). Although our relatively small sample size limits the power of this analysis, we find three significant correlations and two near-significant correlations after Bonferroni correction for multiple comparisons.

**Fig S12.**
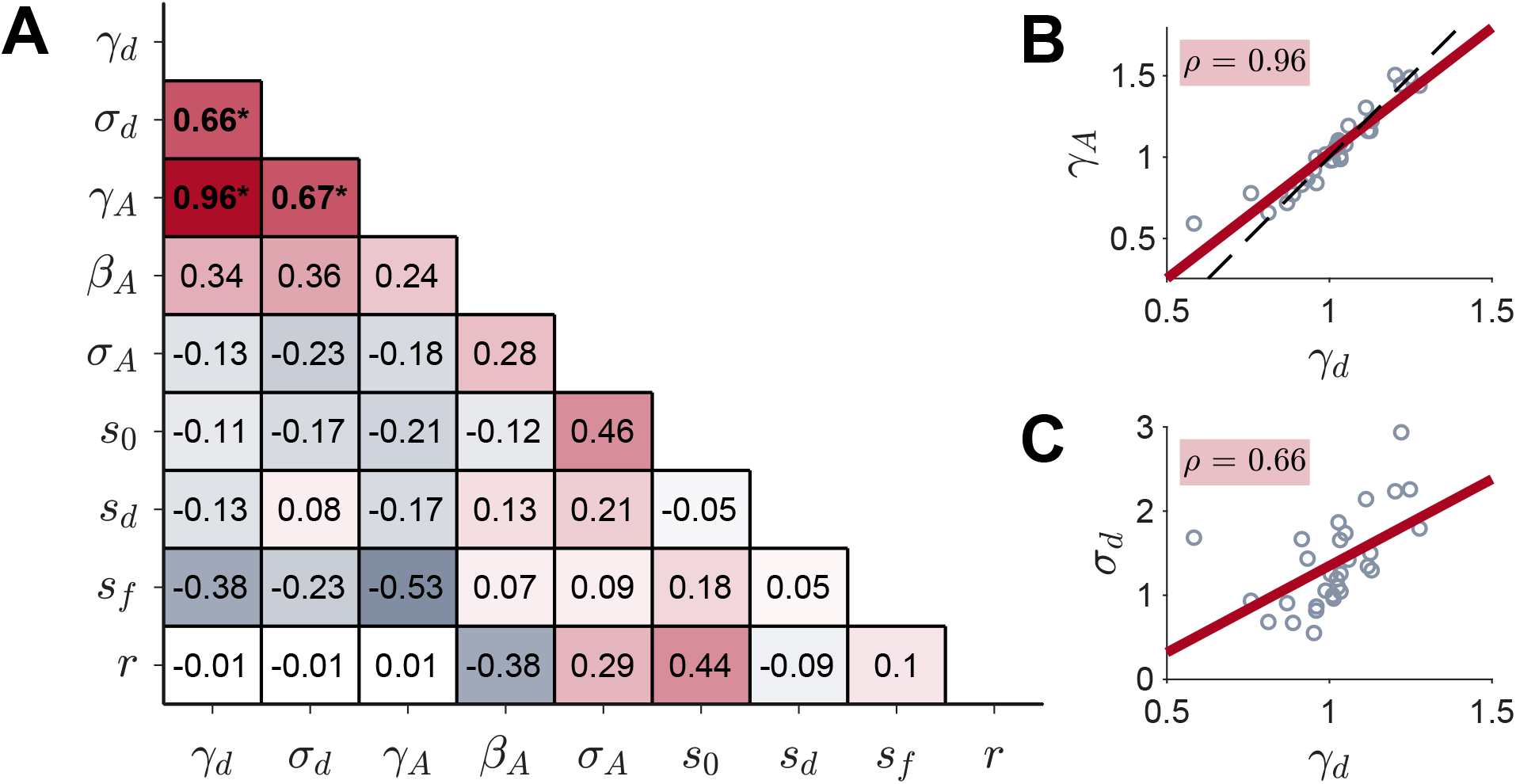
Correlations between parameters in the Hybrid model. (A) Spearman correlation coefficients for all nine parameters. * indicates *p* < 0.05 after Bonferroni correction for multiple comparisons. (B) The correlation between the gain on velocity, *γ*_*d*_, and the gain on the target *γ*_*A*_ is near perfect. The red line corresponds to the linear least squares fit, the black dashed line to the equation *γ*_*A*_ = 2*γ*_*d*_ − 1. (C) *γ*_*d*_ also correlates with the velocity noise, *σ*_*d*_, as does *γ*_*A*_ (not plotted).

The most striking of these correlations is the near-perfect correlation (*r* = 0.96) between the target gain *γ*_*A*_ and velocity gain *γ*_*d*_ (Fig. S12B). As shown by the parameter recovery analysis in Fig. S7, this correlation is not an artefact of the fitting procedure. Instead we believe that this reflects a redundancy in the model whereby the same gain process that contributes to people’s imperfect coding of the target also contributes to imperfect coding of velocity. Intriguingly, the correlation in Fig. S12B is almost perfectly described by the equation

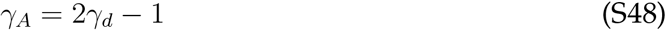

which is the dashed black line in Fig. S12B.

This relationship can cause displacement biases of target location in the direction of motion, which is a phenomenon in perception called *representational momentum* [57–59]. This displacement, characterized as a memory bias, is directly influenced by velocity and has a linear relationship for small changes in velocity [60, 61]. Thus, we speculate that this linear relationship between this deviation from perfect gain (i.e. gain = 1) in memory and velocity is the derivative equivalent to representational momentum. However, exactly why the slope of this relationship should be 2 is a mystery to us at this stage.

The other significant (and near significant) correlations also involve *γ*_*d*_ and *γ*_*A*_ with the velocity noise *σ*_*d*_ and the participant’s estimate of feedback noise *s*_*f*_. Because *γ*_*d*_ and *γ*_*A*_ are so tightly coupled, it is not surprising that their correlations with *σ*_*d*_ and *s*_*f*_ are almost identical and we focus only on the correlations with *γ*_*d*_ in Fig. S12.

For velocity noise, a positive correlation with *γ*_*d*_ (Fig. S12C) can be understood if the noise in the velocity estimate occurs *before* the gain is applied. This is consistent with modifying the equation for the noisy velocity to be

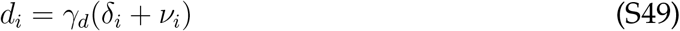

where the standard deviation of the noise is *k*, which relates to the standard deviation of the noise in the original model as *σ*_*d*_ = *γ*_*d*_ × *k*.

Finally, while asserting the null comes with serious caveats with such a small sample size, we note one correlation that was not significant. In particular, we note that *s*_*d*_ and *σ*_*d*_ are only weakly correlated. Perfect Bayesian inference would have these equal, as participants use their estimate of their own velocity noise to optimally integrate feedback. If our model is correct, then this suggests that participants may not have a good estimate of their own path integration noise.

### 5 Model fit for all subjects

**Fig S13.**
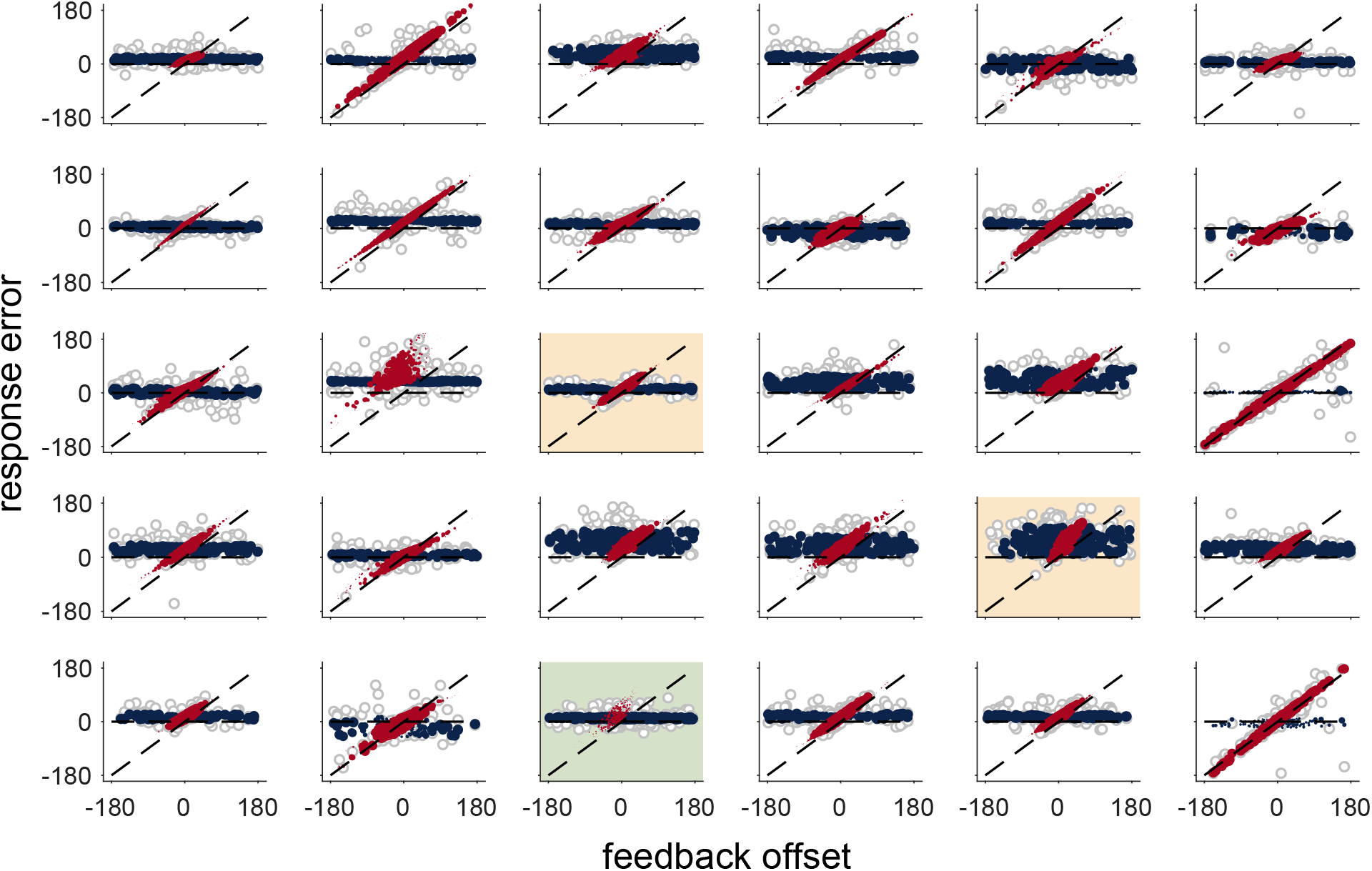
Comparison between data and model for all participants.

### 6 Confidence Rating Correlations

At the end of each trial, participants rated their confidence by adjusting the angle *ς* in Fig. 1D. We found no correlation between any participant’s confidence rating and their angle error (Fig. S14), which is consistent with previous work showing that people are relatively bad at judging their own errors [62, 63]. Interestingly some participant’s confidence rating does correlate with target angle Fig. S15. In other words, these participants feel less and less confidant as they continue rotating. One possible explanation that these participants not only estimating target location but rather they are calculating full posterior distribution [64]. Indeed some of these participants show a significant correlation with the posterior variance calculated from the Hybrid model Fig. S16. However, with the larger individual differences, it is hard to make an exact conclusion.

**Fig S14.**
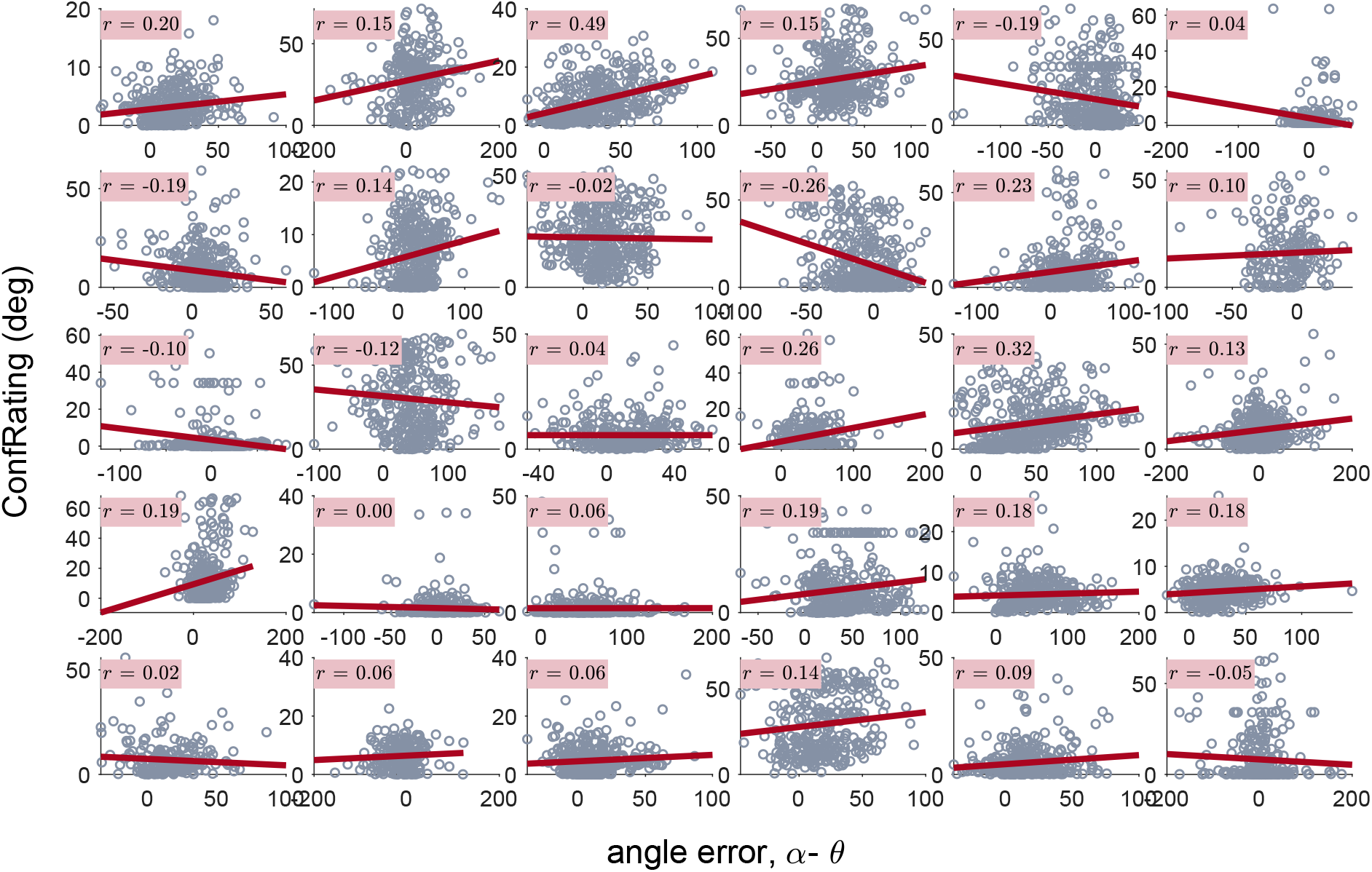
Subject’s confidence rating plotted against their angle error

**Fig S15.**
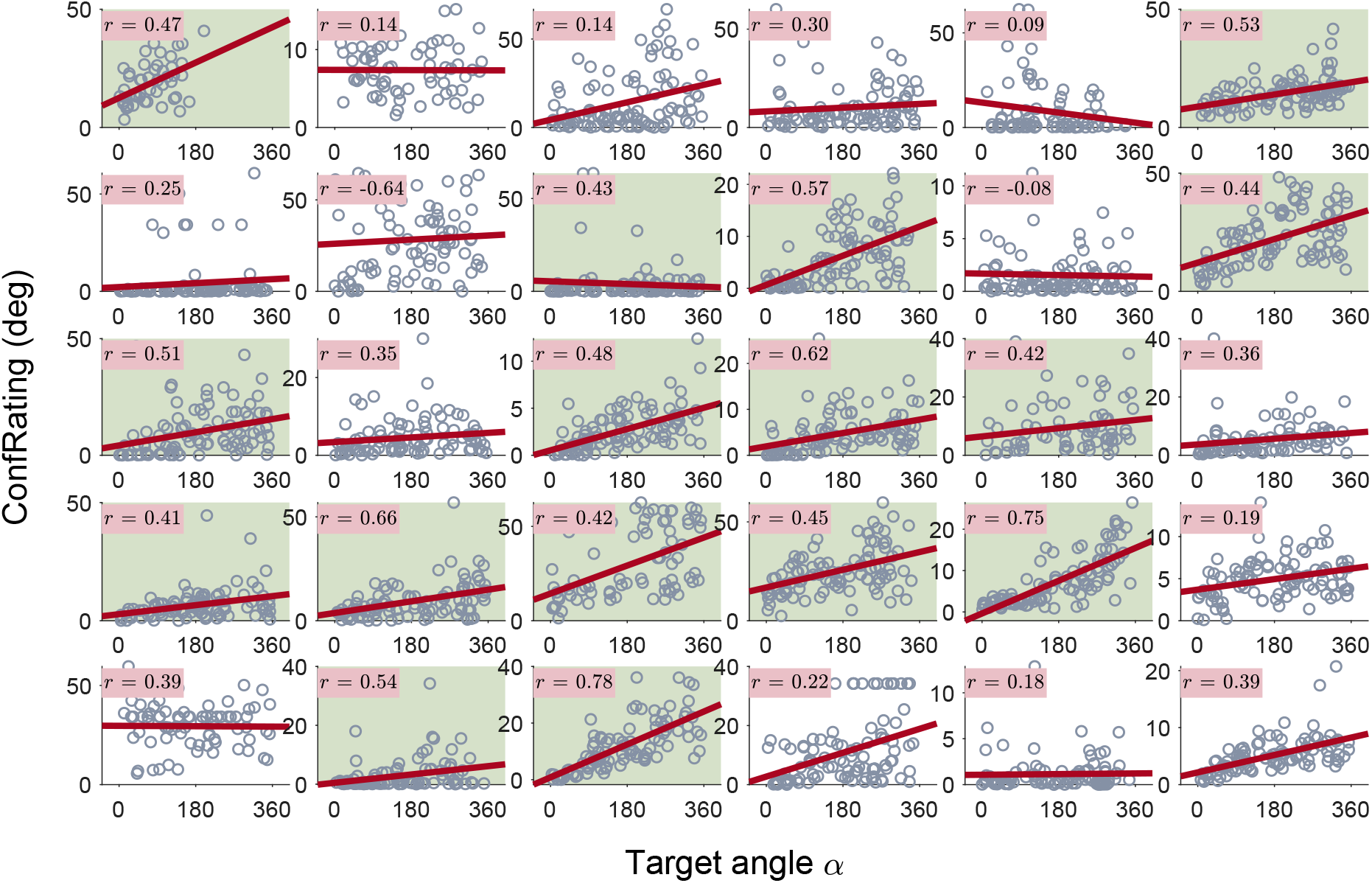
Subject’s confidence rating plotted against target location. Green: *r* >≥ 0.4

**Fig S16.**
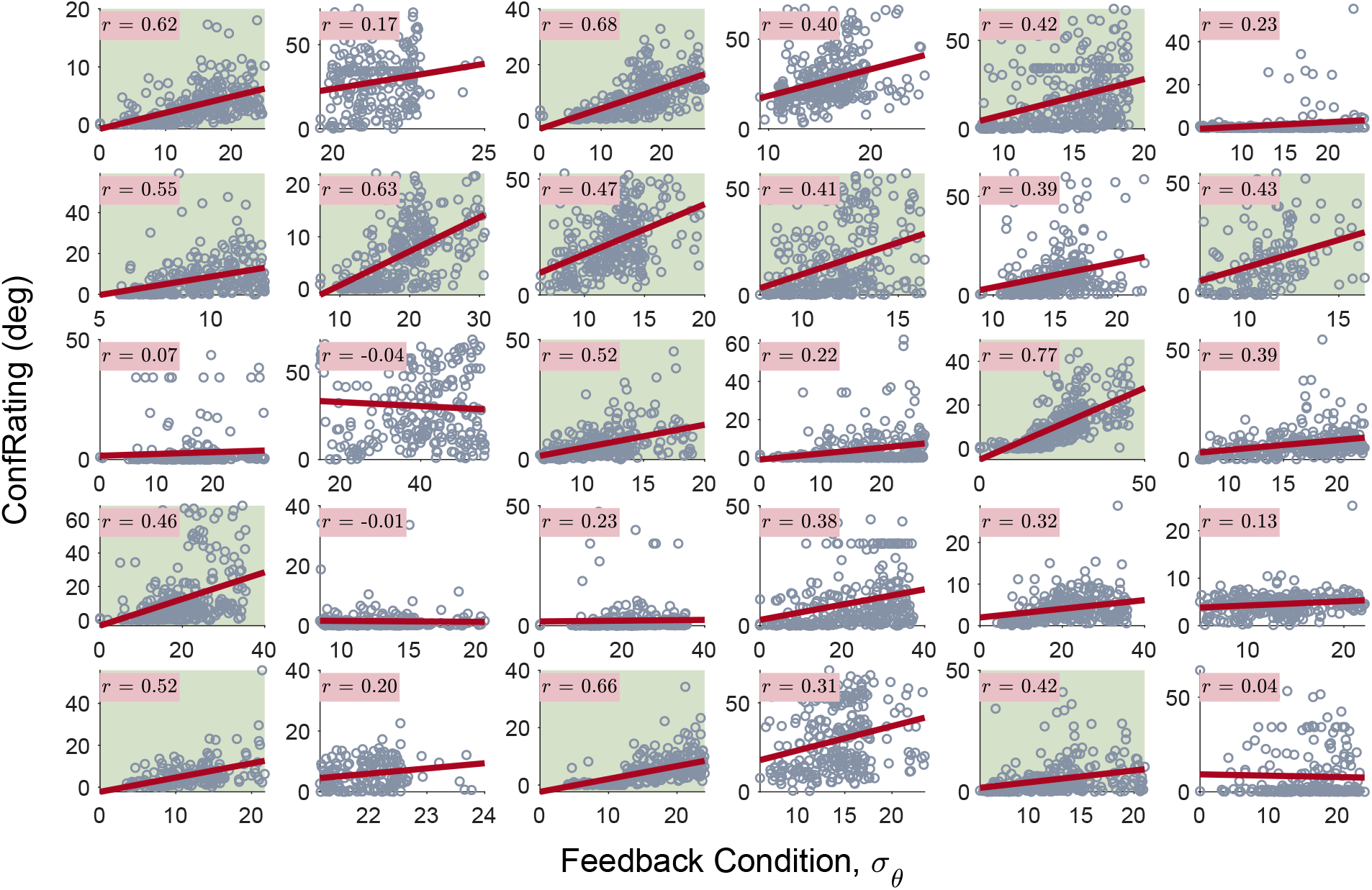
Subject’s confidence rating plotted against the posterior variance Eq. 13 and S36

